# General, orders-of-magnitude faster whole-genome analysis with genotype representation graphs

**DOI:** 10.64898/2026.04.10.717786

**Authors:** Drew DeHaas, Chris Adonizio, Ziqing Pan, Xinzhu Wei

## Abstract

Genotype Representation Graphs (GRGs) are a biologically-motivated, graph-based representation that supports computation directly on the losslessly compressed genotypes. We present GRG v2, which reduces GRG construction time by 10-20× over v1, while halving the disk and RAM footprint of the resulting files. On the UK Biobank (N=490,541, M=706,556,181), GRG v2 files are 25× smaller than .*vcf.gz* and 8× smaller than PGEN. We also introduce *grapp*, a library with GRG-based pipelines for variant and sample filtering, genome-wide association studies (GWAS) with covariates, principal component analysis (PCA), and data export. *grapp* provides linear operators compatible with *numpy* and *scipy*, enabling matrix multiplication against the standardized genotype matrix, the linkage disequilibrium matrix, and the genetic relatedness matrix all via GRG. Our *scipy*-based PCA runs 51–492× faster than existing methods, enabling a leave-one-chromosome-out approach to GWAS covariate construction, illustrating the scalability and methodological flexibility GRG achieves.

## Introduction

Whole-genome next generation sequencing (WGS) biobank-scale cohorts have fundamentally changed the scale of data available for population and statistical genetic analyses. The recently phased UK Biobank WGS dataset contains more than 700 million variants across nearly half a million individuals. In contrast, the previous whole-genome imputed dataset contains ∼96 million variants. At these scales, popular tabular formats like .*vcf.gz* (gzipped variant call format ^1^) and BED (*PLINK1* ^2^)^3^ struggle to support the demand for computation and storage. Computing simple statistics like allele frequency or filtering a dataset for quality control (QC) can take hours to days. More complex calculations, such as those for statistical and population genetics, can be computationally infeasible unless the variant set has been aggressively filtered.

As real datasets increase in size, so does the need for correspondingly large simulated datasets, for validating the performance and accuracy of computational methods. Two popular methods for such simulations are msprime ^4^ and SLiM ^5^, both of which can be used for realistic human (or other organism) simulations via *stdpopsim* ^6^. These simulated datasets are naturally stored as Ancestral Recombination Graphs (ARGs), which compactly encode the genetic history and variant data of each sample. Genotype data embedded in an ARG stored in the commonly used *tskit* tree-sequence format ^7^ needs to be exported to .*vcf.gz* for use with other tools, but this export is slow, and the resulting tabular format is computationally inefficient.

Principal Component Analysis (PCA), a commonly used unsupervised method for capturing the ancestry of a set of genomes, exemplifies this computational challenge. The largest *k* principal components (PCs) are routinely used as covariates during GWAS to control for population stratification ^8,9^, as a way to visualize geographic ancestry ^10^, or as inputs for positive selection scans ^11,12^. Over the last decade, substantial methodological work has gone into accelerating PCA on genetic data, based on random projection ^11^, iterative eigen decomposition ^13,14^, and optimization of a probabilistic model ^12^. Typically, PCA is run on hundreds of thousands of variants at most, through LD pruning and minor allele frequency filtering ^15^, a workaround that is based on both methodological and computational rationales. However, even with LD-pruned or filtered variants, biobank-scale genotype matrices cannot fit in a reasonable amount of RAM, hence existing PCA methods have to load/use a portion of the genotype matrix at a time to get around the RAM constraint. Many other analyses that rely on information from the whole-chromosome or whole-genome also bear the same constraints.

More recently, alternative genotype representations have been proposed as a possible solution ^16–18^. Notably, Genotype Representation Graphs (GRGs) ^19^ bridge the gap between ARGs - which are a compact, hierarchical representation - and tabular formats, where most real data lives. A GRG can be very efficiently constructed from either tabular formats like .*vcf.gz* and IGD ^20^, or from an ARG in *tskit* format (e.g., simulated data). Without using traditional compression, GRG compresses genomic data by exploiting the implicit hierarchy in the data: the shared ancestry from the samples in the dataset. Version 1 (v1) of GRG was able to store hundreds of millions of variants for millions of haplotypes in files 10-20x smaller than .*vcf.gz*, while supporting graph traversals that can speed up filtering and computation tasks by multiple orders of magnitude. For example, phenotype simulation via GRG can be hundreds of times faster than via an ARG-based method ^21^. However, GRG v1 had several practical limitations: graph construction did not account for variation in diversity along the genome, loading a graph from disk was slow and RAM-intensive, and there was no API for matrix multiplication, which is an operation underlying most statistical genetics methods. Equally importantly, GRG lacked a software ecosystem for broad adoption. The recently released *snputils* library ^22^ supports GRG as one of the four input formats (the others being VCF, BED, and BGEN^3^), but reads GRG into a tabular representation, and so does not benefit from the graph’s structural advantages during analysis.

Here we describe two contributions that together make GRG a practical basis for biobank-scale population and statistical genetics. First, we introduce GRG v2, a substantially improved version of the format and construction algorithm. GRG v2 reduces construction time by 10-20×, halves the disk and RAM footprint of the resulting GRG (resulting in files 20-40 times smaller than .*vcf.gz*), and improves load time by more than 20× relative to v1. Constructing a GRG for the full UK Biobank WGS dataset (490,541 individuals, 706,556,181 variants) costs about 88 GBP (from IGD) or 242 GBP (from .*vcf.gz*, which includes the cost to convert to IGD) and produces files more than 8 times smaller than *PLINK2*’s PGEN ^23^ format. Second, and more importantly, we introduce *grapp*, a pure Python library and command-line tool that exploits the computational advantages of GRG for both routine large-scale analyses and new method development. *grapp* implements efficient GWAS with covariates, PCA, export from GRG to IGD, filtering by samples or variants, and an extensible library of linear algebra operations that are compatible with the *scipy* ^24^ and *numpy* ^25^ ecosystem. We demonstrate the practical impact of this framework through applying the new GRG functionalities to the UKB WGS dataset.

## Results

### Overview of Genotype Representation Graphs

A GRG losslessly encodes genotypes in a directed acyclic graph that makes up a multi-tree, where variants and samples are both mapped to nodes in the graph (Figure 1ab). Samples *s*_*i*_ are the leaves of the multi-tree, and the variants *v*_*j*_ (called “Mutations” in the GRG libraries) can be mapped to any node in the graph. All samples reachable from the node for variant *v*_*j*_ contain *v*_*j*_ in their genome. Conversely, all variant nodes reachable from a given sample *s*_*i*_ make up the haplotype for *s*_*i*_. Therefore, subsets of the genotype data based on a list of variants or samples can be obtained from a GRG by collecting all reachable nodes via downward traversal (from the desired variants) or via upward traversal (from the desired samples). For example, in Figure 1b the node for variant *v*_*1*_ reaches samples *s*_*1*_, *s*_*2*_, *s*_*3*_, indicating that *v*_*1*_ is present in their haplotypes. Traversing upward from *s*_*3*_ reaches *v*_*1*_, *v*_*2*_, and *v*_*3*_, meaning those variants are all present in *s*_*3*_’s haplotype.

**Figure 1:**
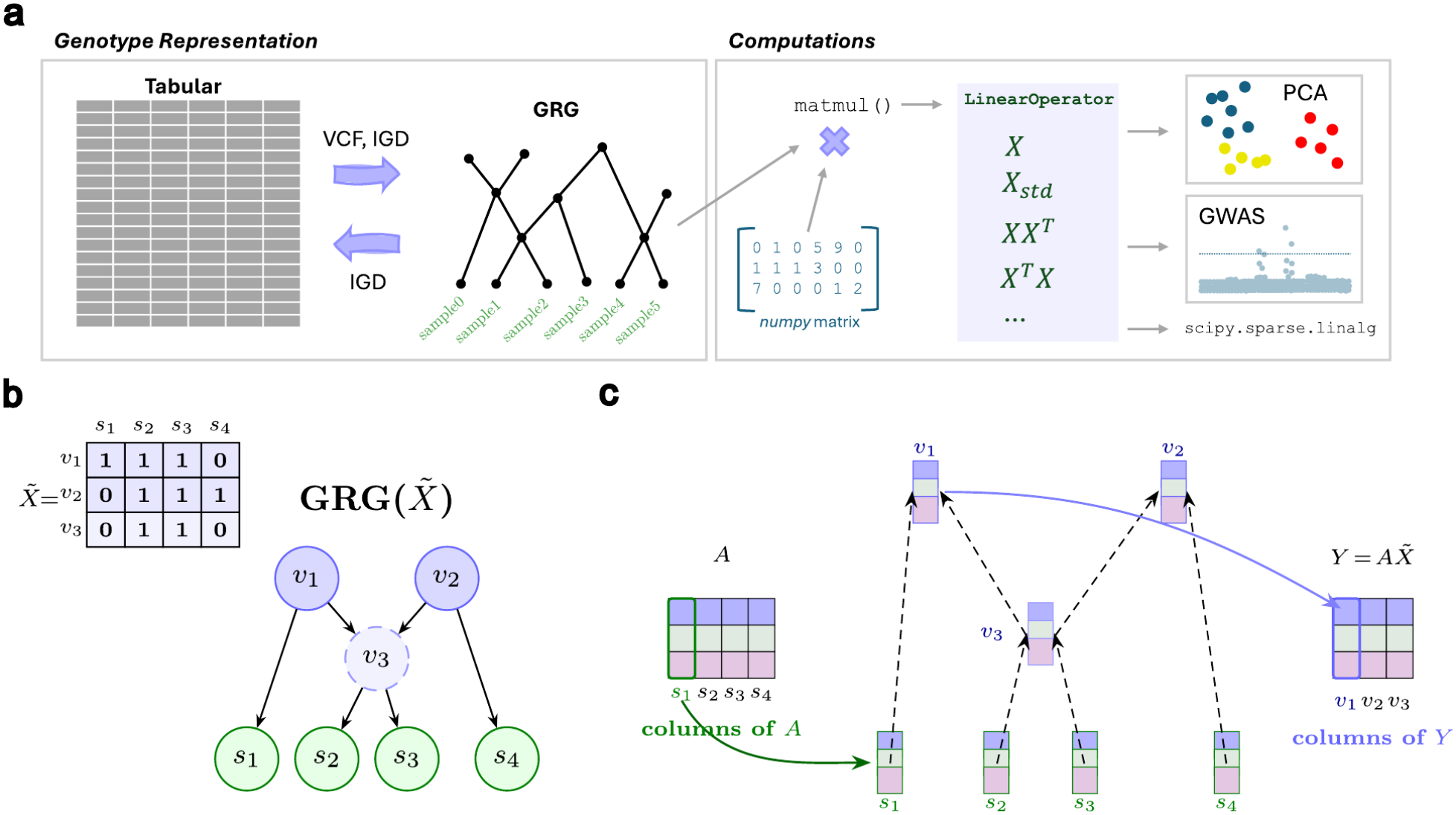
Graph-based matrix representation and multiplication. **a**. A GRG represents the same genotype matrix as tabular formats, and can be constructed from VCF or IGD. Matrix multiplication against GRG integrates with *numpy* and *scipy* to enable statistical genetics calculations. **b**. An example of the transpose of a small haploid genotype matrix 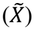 and a corresponding GRG. Samples are *s*_i_, and variants are *v*_j_. **c**. Diagram illustrating matrix multiplication 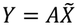 where *A* is a dense matrix and 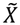 is the genotype matrix, which is encoded as a GRG. This is an upward traversal where each node stores a column of values which is summed element-wise from the child nodes. The sample node values are initialized from the corresponding columns of *A*.

A GRG speeds up computation by representing repeated intermediate results with shared nodes, so each can be computed once and reused during graph traversal. In addition to the typical graph traversals (topological order, depth-first search, breadth-first search), GRG v2 supports matrix multiplication as a graph operation (Figure 1c). If *X* represents the *N* × *M* genotype matrix for a GRG *G*, then the product *AX* can be obtained through an upward topological traversal of *G* by setting the values of the sample nodes to *A* and then setting each node’s value to the sum of its child nodes values. Similarly, the product *AX*^*T*^ can be obtained by a downward topological traversal. The equivalence of GRG-based matrix multiplication to standard matrix multiplication has been demonstrated in Syam et al.^21^. Given a *K* × *N* matrix *A* being multiplied against a genotype matrix, the traditional matrix multiplication is *O*(*KNM*) whereas the GRG-based algorithm is *O*(*K*|*G*|) where |*G*| is the number of edges in the graph. A naive GRG can be seen as a bipartite graph with *M* variant nodes, *N* sample nodes, and *C*_*i*_ allele copies for variant *i*, such that |*G*| = ∑_*i*_ *C*_*i*_; whereas the GRG construction algorithm builds shared hierarchy to ensure that |*G*| << ∑_*i*_ *C*_*i*_.

### Fast GRG construction for 490,541 genomes and 706,556,181 variants

The primary step of GRG v2 construction (see Methods) is called *Build*, and it is applied to short (typically 50-100 KBP) regions of the genome to build multi-trees, which are then combined into a single larger GRG. The *Build* algorithm performs neighbor joining of all haplotypes recursively, using Hamming distance as a metric. The primary improvement to *Build* in GRG v2 is a more compact and efficient haplotype representation (Supplemental Figure S1) that allows lossless representation of all haplotypes (for a genomic region) in a small RAM footprint. In GRG v1, the haplotype representation used during graph construction was lossy, which necessitated a process with two major steps: the *BuildShape* step that creates a graph without mutations, and a second time-consuming step *MapMutations* that modified the graph as mutations were added. In GRG v2, *MapMutations* is no longer necessary, as the node-to-mutation mapping is now maintained during *Build*. The *Build* function creates the tree by iteratively joining nodes, starting with just the sample nodes, and the shared haplotype (intersection of variants) from the joined children nodes is stored at their parent node. In addition to the shared parent node (which is also created by *BuildShape*), the *Build step* creates an extra parent node for each child, a “left” and a “right” node that store the left and right set differences of the joined children (Figure 2a), which means that the tree (now a simple multi-tree) losslessly stores the mutation-to-sample relationship. We then collect all the nodes containing each mutation, and create a single mutation node with edges to those nodes (Figure 2b). Furthermore, we introduced an improved edge list encoding (see “Improved Graph Construction” in Methods) in GRG v2, which reduced both file size and RAM footprint.

**Figure 2:**
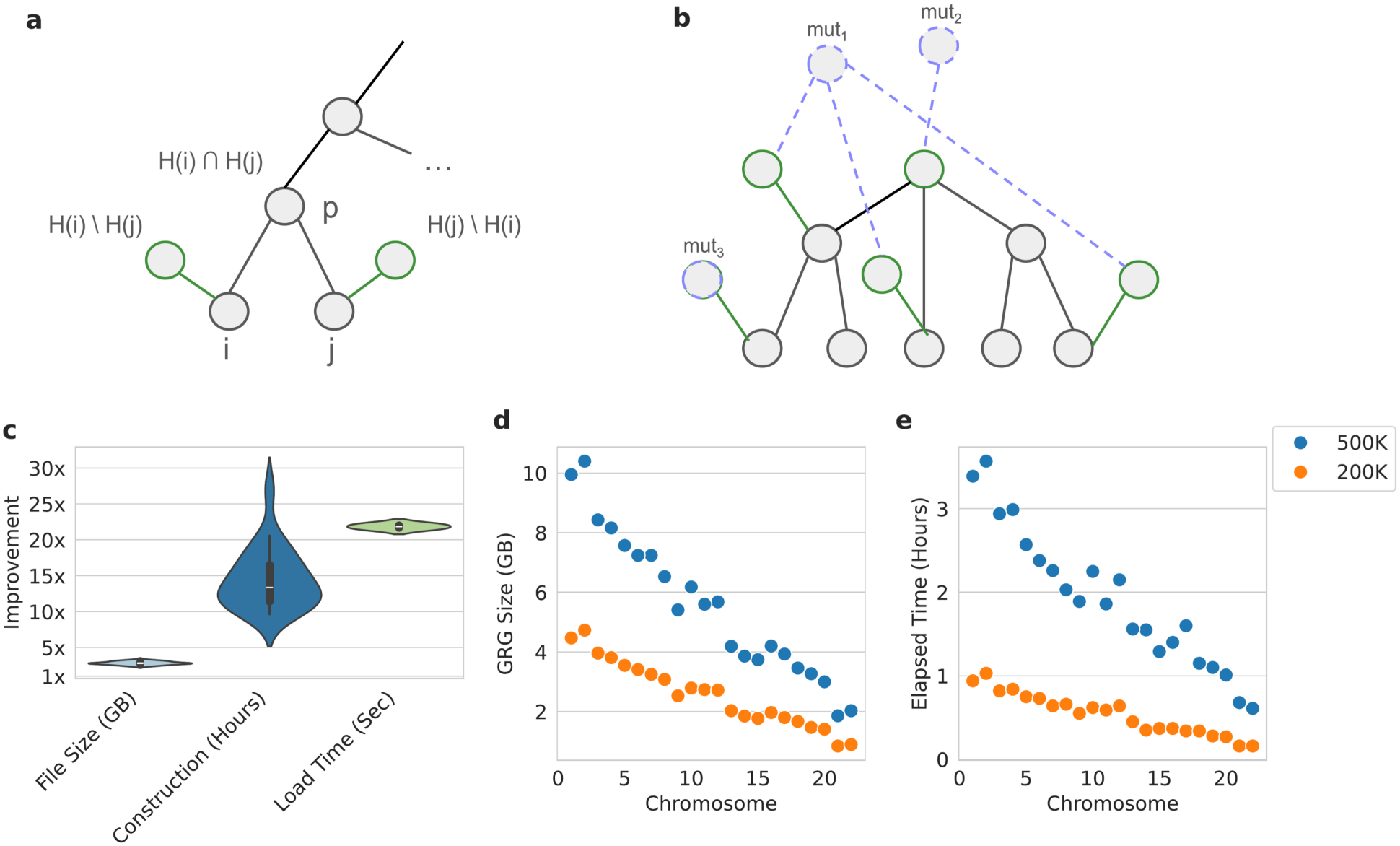
Improved GRG results on UK Biobank. **a**. Illustration of GRG *Build* algorithm, where H(*i*) is the haplotype (i.e., set of variants) shared by all samples beneath node *i*. In this example, node *p* captures the common set of variants between node *i* and *j*, where H(*i*)∩H(*j*) represents the set intersect. The green nodes become roots and capture the “residual haplotypes” that do not reach the parent of nodes *i* and *j*, or the set differences (H(*i*)\H(*j*), and H(*j*)\H(*i*)). Because {H(*i*)\H(*j*)}∪{H(*i*)∩H(*j*)}= H(*i*), the graph emitted from *Build* contains all the haplotype to variant information. **b**. After the tree is constructed, all root nodes (green) are connected to the relevant mutation nodes (light blue) for the haplotypes they capture. **c**. Improvement of GRG v2 over v1 on the UK Biobank 200,000 phased individuals WGS dataset, reflecting how much smaller and faster our improved GRG construction and file format are. **d**. GRG file sizes for both the 490,541 (500k) and 200,011 (200k) phased individuals WGS datasets. **e**. Elapsed time for constructing GRGs from IGD with 64 threads (500K dataset) and 70 threads (200K dataset).

Collectively, these changes resulted in a suite of improvements over GRG v1 (Fig. 2c, using 200,000 UKB WGS), including halving the size of GRG, 10-20× faster construction, and more than 20× faster load time (263.7 vs. 11.9 seconds, for chromosome 13). Total construction cost of 87.71GBP (Supplementary Table S3) is from a mixture of spot and on-demand instances; larger chromosomes used on-demand jobs. We benchmarked GRG v2 construction on both the 200,000 and 500,000 individual UKB datasets (Figure 2d-e). The file size on the UKB datasets appears approximately linear in the number of samples, whereas the scaling for simulated datasets is likely sub-linear (Supplemental Figure S2). Noise or other complexities in real data (not captured by the simulation model) are likely the cause of this scaling difference. Maximum RAM usage while constructing 500,000 individual GRGs varied from 92.5GB (chromosome 22) to 241.8GB (chromosome 2) (Supplemental Table S1). The user can reduce RAM usage during construction by requesting the genome be split into more parts (the *-p* option), at the cost of a potentially larger GRG file in the end.

Comparing with *PLINK2*’s PGEN format, we see that GRG is more than 8× smaller (Table 1) while requiring similar time to construct from VCF files. IGD files, a tabular format similar to .*vcf.gz* in simplicity but stored in binary (not text) form, offers a substantial speed advantage for constructing biobank-scale GRGs (almost 3× faster than .*vcf.gz* for UKB chr22).

**Table 1:** Constructing a GRG vs. constructing PGEN.

| **Chrom.<sup>1</sup>** | **Format** | **File Size (GB)** | **Threads** | **Input** | **Minutes Elapsed** | **Minutes CPU<sup>2</sup>** |
|---|---|---|---|---|---|---|
| 22 | PGEN | 17.40 | 2 | .vcf.gz | 1247.62 | 2495.24 |
| 22 | PGEN | 17.40 | 70 | .vcf.gz | 335.40 | 23478.00 |
| 22 | GRG | 1.98 | 70 | .vcf.gz | 100.72 | 7050.40 |
| 22 | GRG | 2.03 | 70 | IGD | 36.60 | 2562.00 |

<sup>1</sup> UK Biobank chromosome 22 phased WGS data (490,541 individuals).
<sup>2</sup> This is elapsed time multiplied with the number of threads, not the operating system’s measurement of CPU usage.

Once data is in GRG format, it can be filtered very efficiently by variants or samples (Supplemental Figure S3). However, newly generated or expanded datasets are usually stored in a tabular format like .*vcf.gz*. Thus, the time, cost, and cloud node requirements for constructing a more efficient format like GRG should still be a consideration.

### PCA on 137 million UK Biobank variants

Over the past decade, PCA has become a standard step in population and statistical genetic analyses. With WGS data, the total number of variants across all autosomes can be in the hundreds of millions, hence existing methods have to deal with large genotype matrices that don’t fit in a reasonable amount of RAM, through only loading/using a portion of the matrix at a time. In contrast, the GRG representation is substantially smaller than the matrix representation (dense or sparse), and can fit the entire graph into RAM, while also needing fewer floating point operations (on the order of O(|*G*|)) to perform the multiplications, both of which offer a substantial speedup over existing methods and enable use on WGS data.

*grapp* implements two GRG-based PCA methods: one based on iteratively solving the eigen decomposition via scipy (referred to as “GRG” PCA), and GRG ProPCA (based on the ProPCA algorithm ^12^) (Figure 3ab). We compared the runtime performance of GRG to compute the top 10 PCs with two other PCA methods: *PLINK*2 ^23^ with the “approx” option, which uses a randomized algorithm akin to FastPCA^11^, and *FlashPCA2* ^13^ (Figure 3ab). We used simulated datasets based on the *stdpopsim* OutOfAfrica_4J17 model ^26^, sampling equally from all four populations. PCA jobs were limited to up to 240 hours. Our *PLINK*2 tests use the PGEN format and *FlashPCA2* uses BED format. Since most existing methods are intended to work on smaller variant sets, we also tested these methods after filtering the datasets to exclude variants with derived allele frequency < *0.05* (Figure 3cd). We allotted up to 128GB of RAM for each tool, and gave *FlashPCA2* a block size of 100GB (in order to speed up its analysis). *PLINK*2 and *FlashPCA2* were both allowed 25 threads (though *FlashPCA2* only made use of 1), and GRG’s PCA is single threaded per-graph (this experiment used a single graph). In both the filtered and unfiltered dataset, GRG is faster than the other methods and uses less RAM. Both GRG methods are multiple orders of magnitude faster than the other methods on the unfiltered dataset (51-492 times faster than PGEN), and the eigen-decomposition based GRG method uses the least RAM of all methods in both datasets (PGEN uses 20-27 times more RAM). For example, with the unfiltered 500,000 individuals dataset, GRG PCA (single threaded) took 14.3 minutes (3.3GB RAM), whereas the fastest non-GRG method (*PLINK2* with 25 threads) took 39.1 hours (117GB RAM).

**Figure 3:**
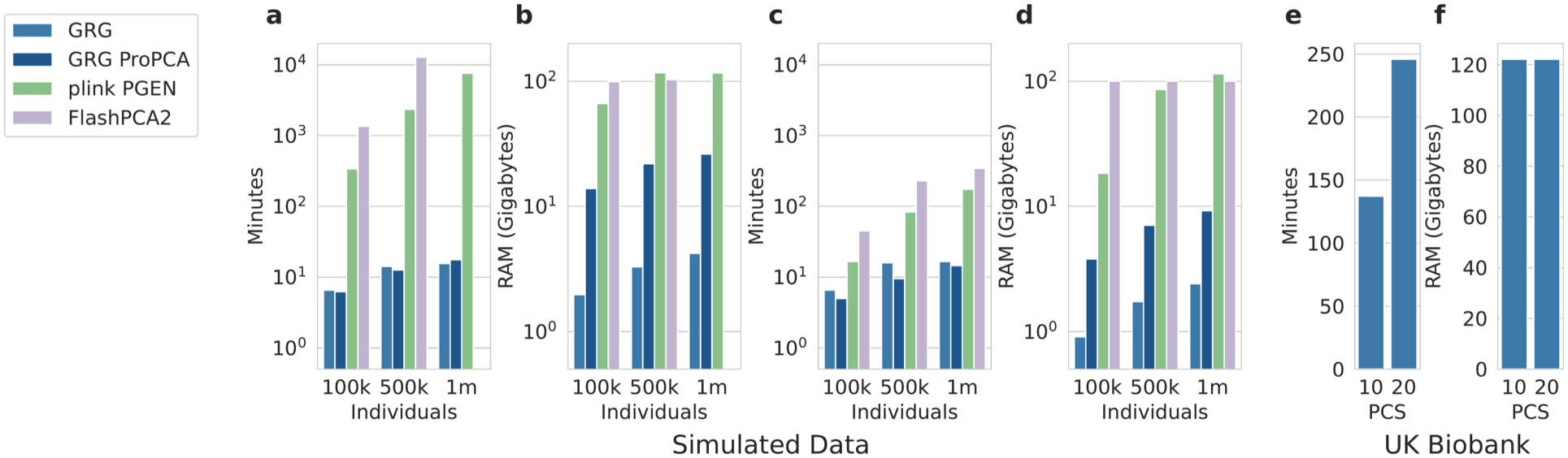
PCA runtime performance and example applications. **a**. Comparing times to obtain top 10 PCs on unfiltered simulated data of different sample sizes. Datasets contain between 14.5 and 23.7 million variants. **b**. The same tests as panel **a**, but showing RAM usage. **c**. Comparing times to obtain top 10 PCs on simulated data filtered such that derived allele frequency >0.05. Datasets contain about 440,000 variants. **d**. The same tests as panel **c**, but showing RAM usage. For panels **a-d**, the y-axis is log scale. **e**. Time to run PCA on the UK Biobank dataset with 490,541 individuals and 137,116,837 variants over 22 autosomes with 22 threads. **f**. Same as panel **e**, except RAM usage.

We applied GRG PCA (*k* = 10 and *k* = 20) on the recently phased 490,541 individuals (WGS) in the UK Biobank. We first filtered out ultra-rare variants (allele count < 20), then ran GRG PCA on all the remaining variants in autosomes (137,116,837 variants), with one GRG per chromosome, which required ∼122GB of RAM and took 2.3 (*k* = 10) to 4.1 (*k* = 20) hours (using 22 threads, one per GRG). Overall, the largest 3 PCs across populations were broadly consistent with the “genetic principal components” field provided in the UK Biobank (field 22009), though the GRG PCs were extracted from a dataset with more than 900 times the number of variants (Supplemental Figure S4).

### Graph-based GWAS with covariates

In this work, we extended this graph-based GWAS, previously described in DeHaas, et al., 2025^19^, to support covariates. Unlike PCA, which requires repeated matrix multiplications against the genotype matrix until convergence, GWAS can be done in a fixed number of multiplications (3 when covariates are used, 2 when they are not), and can also be done variant-by-variant, which makes it more amenable to computation with tabular formats.We compared computation time (Figure 4a), RAM usage (Figure 4b), and p-values (Figure 4c) between *grapp* and *PLINK*, running GWAS with the top 10 PCs as covariates. Because the entire GRG is loaded by *grapp* and *PLINK* is able to load data variant-by-variant, *grapp* is expected to use more RAM. However, in these datasets, the RAM difference is only by a factor of 2-4×. *grapp* (one thread, since there is one GRG) is faster than single-threaded *PLINK* on all datasets, and as the number of individuals in the dataset increases, it becomes faster than *PLINK* with 25 threads. *grapp*’s p-values correlate very strongly with *PLINK*’s (for 99.8% of the tested variants, the p-values differ by less than 1%), with the differences due to the handling of multi-allelic sites (*plink* appears to model the alleles jointly, *grapp* independently).

**Figure 4:**
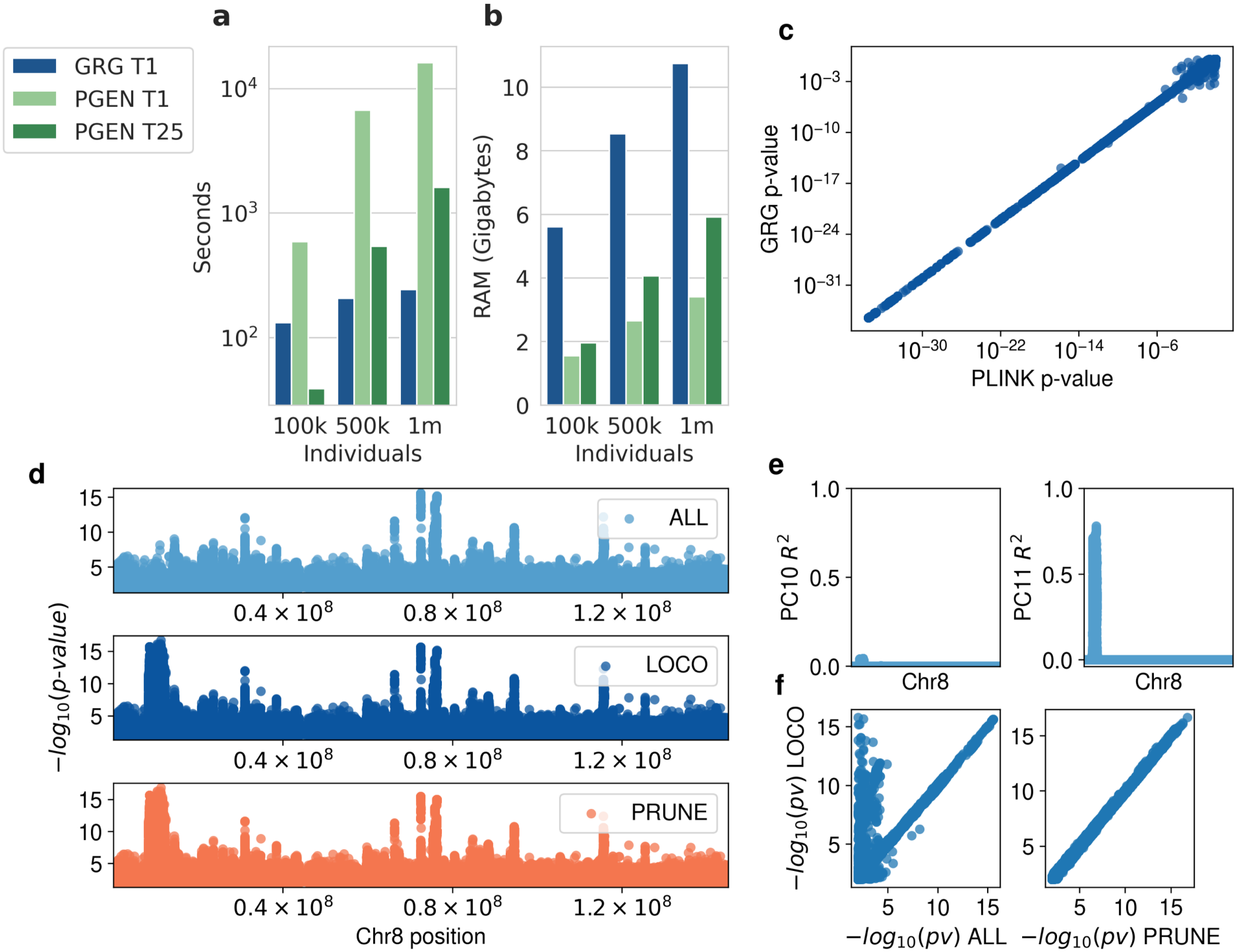
GWAS with covariates. **a**. Time to perform the same GWAS (with 10 PCs as covariates) on simulated data for GRG and *PLINK*2 PGEN. GRG uses 1 thread (GRG T1), *PLINK2* was given 1 (PGEN T1) or 25 (PGEN T25) threads. **b**. Same as panel **a**, but RAM usage. **c**. *PLINK*2 vs. *grapp* p-values for the same GWAS (PCA covariates) on simulated data and phenotype. **d**. GWAS p-values for body mass index (BMI) on UK Biobank chromosome 8, unrelated white British individuals, covariates are the top 20 PCs and sex. Each Manhattan plot uses PCs from a different PCA method: PCA on all autosomes (ALL), leave-one-chromosome out (LOCO) PCA, or the LD-pruned PCA (PRUNE, from UK Biobank field 22009). **e**. Correlation between PC10 (PC11) and the chromosome 8 genotypes, with PCs computed from PCA on all autosomes (ALL). **f**. p-values from GWAS using covariate PCs from LOCO (y-axis), vs. from ALL (x-axis, left) or PRUNE (x-axis, right).

A practical issue with using PCs as covariates is that they capture more than just global ancestry or within-population stratification: strong signals of linkage disequilibrium (LD) and relatedness can also show up in PCs. Grinde, et al.^27^ demonstrated that effect sizes for (strongly) causal variants can be biased due to LD captured by PCA. Although, “LD pruning” is a standard procedure to control for such biases, by removing all nearby variants that are highly correlated with each other, the optimal parameters (i.e., window-size and *r*^*2*^ threshold ) for LD pruning vary across datasets and might require additional parameter tuning ^27^. Given that GRG-based PCA is computationally cheap, even on variant-dense datasets, we instead propose a different method to avoid LD in the PCs used for GWAS: the leave-one-chromosome-out (LOCO) approach. For each autosome *c*, we perform PCA on all the autosomes together, except for *c*, and the resulting PCs become covariates for the GWAS performed over *c*. Unlike LD pruning, LOCO has no parameters, and does not modify the dataset, both of which aid reproducibility and consistency. Using this approach, no within-chromosome LD within *c* is present in the covariates, as shown in the manhattan plot for GWAS of BMI (Figure 4d) which shows an LD-related effect on variant p-values when using all unpruned autosomes for PCA (“ALL”), compared to the LOCO and LD-pruned PCs (the latter are released by the UK Biobank, as field 22009). The p-values from the LOCO-based GWAS are very consistent with the p-values from the LD-pruned-based GWAS, and deviate substantially from the naive use of all autosomes for PCA (Figure 4f).

We also directly examined the correlation between genotypes and PCs by performing a regression between each variant and each PC, and reporting the coefficient of determination (Supplemental Figure S5). For example, the undesired p-value effect on the BMI GWAS with chromosome 8 (Figure 4d) is likely explained by the high genotype-PC correlation in PC10 and PC11 when the PCA method does not control for LD (Figure 4e). Interestingly, for the chromosomes we examined (5, 6, 7, 8, 9, and 10) there is a slightly higher genotype-PC *R*^*2*^ with the LD-pruned PCs than with the LOCO PCs (Supplemental Figure S5). In contrast, the PCs from the ALL method show substantial difference from the other two methods, with spikes of genotype/PC correlation within the first 10 PCs for chromosomes 6, 7, and 10, and elevated correlation in some of the later PCs for chromosomes 5 and 8. Depending on the phenotype, the GWAS results may or may not be noticeably affected (Supplemental Figures S9, S11).

### Facilitating efficient workflows with Python Linear Operators

As a Python package with documented APIs and tutorials, *grapp* enables interactive exploration and customized workflows with more flexibility than traditional command-line-based GWAS, PCA, and statistical genetics tools (Figure 5a). While *grapp* provides many pre-implemented features, its integration with *scipy*’s sparse linear algebra libraries allows future methods based on GRG to be developed easily and efficiently. These operators encode not just matrix multiplication against the genotype matrix (*X*) underlying the GRG, but implicit transformations of that genotype matrix. *grapp* supports operators for *X, XX*^*T*^, and *X*^*T*^*X*, where *X* can be the haploid (0, 1-valued) or diploid (0, 1, 2-valued) matrix, and *X* can also be optionally standardized. Additionally, there are multi-GRG versions of these operators, which provide parallel matrix multiplication against multiple GRGs that represent a single matrix (e.g., each autosome is typically a separate GRG file). With these operators, one can write Python code that treats a GRG object like a *numpy* matrix (Figure 5b), while being vastly more efficient (in both RAM and CPU time).

**Figure 5:**
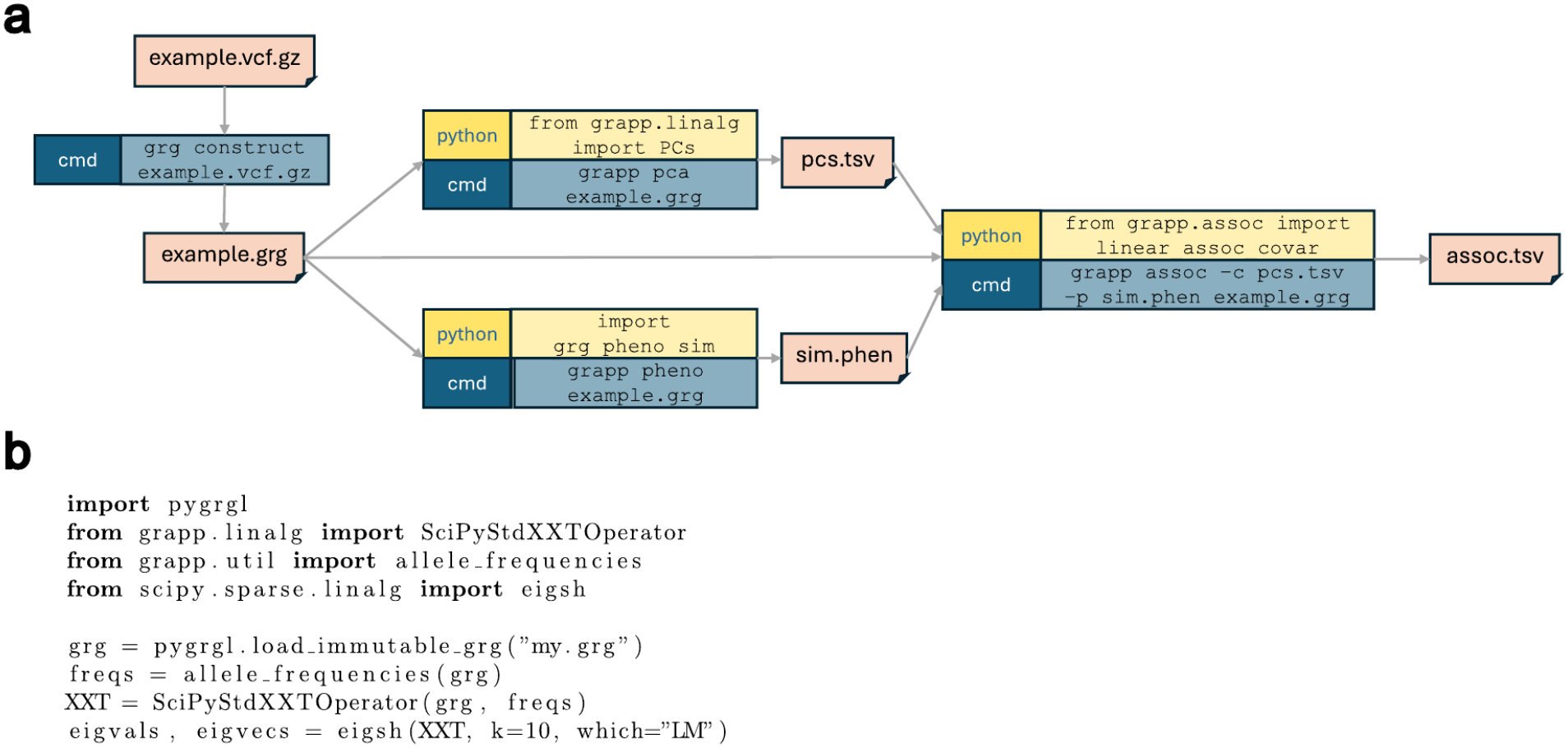
*grapp* usage examples. **a**. Example workflow using a simulated phenotype and PCA covariates as input to GWAS with GRG and *grapp*. **b**. Principal Component Analysis (PCA) implemented with a *grapp* linear operator and *scipy*: eigen decomposition of *XX*^*T*^ on the standardized genotype matrix *X*. Meant to illustrate the simplicity and power of the linear operators; for PCA, users can just use the *grapp.linalg.PCs()* method (which has additional options).

## Discussion

In summary, we introduce several advances to the GRG core library, the improved GRG file format, construction, and *grapp* that together make complex analyses on huge WGS datasets fast, affordable, and easy to implement. Adopting GRG requires minimal up-front investment: constructing a GRG from .*vcf.gz* is now as easy and as fast as converting .*vcf.gz* into PGEN, a routine step in many biobank genetic analyses. Moreover, routine computational analyses such as sample and variant QC, GWAS, and PCA can now be done easily and orders-of-magnitude faster with *grapp*, providing a strong alternative to existing, commonly used tools. Last but not least, the Python linear operators allow new methods to leverage GRG through standard *numpy* and *scipy* interface, opening up possibilities for development, interactive exploration, and customization. Together, these advances position GRG as a practical alternative to tabular formats for biobank-scale population and statistical genetics.

This work opens up possibilities for asking new questions of biobank-scale WGS data and for scaling existing methods to much larger datasets. However, working directly on dense, unfiltered WGS data also exposes statistical issues that filtering had previously moderated. In this study, we explored the interplay between LD and PCA when PCs are used as GWAS covariates. Traditional pipelines apply LD pruning and minor allele frequency thresholds for two distinct reasons: a computational one – keeping the genotype matrix small enough to manipulate – and a statistical one – reducing the influence of correlated and rare variants on downstream methods. When the computational reason disappears, the statistical one becomes more visible. Running PCA on all autosomes simultaneously, without LD pruning, revealed strong local LD signals from chromosome 8 to be captured by PC10 and PC11, biasing the GWAS p-values for variants in that region. These biases can be mitigated by LD pruning, but the optimal window size and *r*^2^ threshold vary across datasets and require tuning, and LD pruning itself becomes computationally expensive as the number of variants increases. The LOCO approach sidesteps the parameter choice entirely by removing the chromosome under analysis from the PCA input, and produces p-values that closely track those from properly tuned LD-pruned PCs. This is practical only because

GRG-based PCA is now cheap enough to run twenty-two times. Notably, although PCA on the full unfiltered variant set is now tractable, ultra rare variants still appear to introduce cross-chromosomal LD-like signals. In UK Biobank data, filtering out variants with MAC<20 prior to LOCO PCA appears sufficient. Other questions about the use of PCs as covariates also deserve revisiting at WGS scale, for example, how many components are needed to capture ancestry and population structure when the input contains hundreds of millions of variants rather than a few hundred thousand. The fast PCA method presented here could facilitate the systematic exploration of these questions on biobank-scale WGS datasets.

Our GRG-based PCA is one example of building an iterative method with GRG. Iterative methods, such as the Arnoldi iteration underlying our PCA implementation, rely on repeated matrix-vector or matrix-matrix products, instead of having to actually manipulate the materialized matrix^28^. *scipy*’s sparse iterative methods accept any object that supports matrix-vector or matrix-matrix multiplication, which is exactly what *grapp*’s linear operators provide. The linear operator interface also provides a nice abstraction for implicit transformations of the genotype matrix, which simplifies the implementation of custom iterative methods, such as *grapp*’s implementation of ProPCA ^12^. This pattern of substituting a GRG-backed operator into existing methods should generalize nicely, as many modern statistical genetics methods are iterative and spend much of their computation on repeated multiplications against the genotype matrix, the LD matrix, or the genetic relatedness matrix. For example, Bayesian variable selection methods ^29^, polygenic score predictions that use the full LD matrix ^30^, and random projection based heritability and genetic correlation estimators ^31,32^, can all be accelerated and scaled to larger datasets via *grapp*’s linear operators. We expect GRG-based linear operators to enable a class of methods that were previously restricted to smaller datasets.

Most of the methods described in this work are about exploiting the compression of a graph to perform mathematical operations. However, the graph itself captures common hierarchies that were formed through ancestral processes, and algorithms that are more graph-centric can potentially exploit this information, similar to ARG-based methods. This suggests a longer-term opportunity: using GRG as a stepping stone toward biobank-scale ARG inference. ARG inference currently remains infeasible for hundreds of millions of variants across half a million individuals ^33,34^. GRG construction, by contrast, scales comfortably to the full UKB WGS dataset. The construction algorithm itself is suggestive of a connection: *Build* recursively merges haplotypes by Hamming distance and stores shared variants at internal nodes, producing a multi-tree whose topology already reflects patterns of shared ancestry, even though it makes no explicit coalescent or recombination assumptions. A natural question is whether the GRG topology, or *Build* itself, can be modified to seed or accelerate ARG inference – for example, by approximating recombination implicitly during graph construction. Realizing this would require characterizing what the GRG topology represents statistically, and developing principled ways to extract substructures with the probabilistic guarantees that ARG-based methods rely on. Such work could move GRG from being an efficient data structure for matrix-based computation to being a useful abstraction for population genetics. Related to this, GRG representation in its current form is suboptimal for inherently sequential tasks, such as finding identity-by-state segments. In the future, it would be an interesting direction to associate genomic intervals with internal nodes and allow a GRG to be traversed based on the order or internal nodes along the genome.

Several limitations and areas for improvements should be noted. While GRG construction is now cheap relative to the cost of downstream analyses, it can require substantial peak RAM (241 GB for the UK Biobank 500k WGS dataset) in order to achieve the best compression, even though analysis-time RAM is much lower. Many of *grapp*’s operations currently operate with a single thread per GRG, where each GRG typically represents a chromosome; while this is already much faster than multi-threaded alternatives in many cases, there is room for improvement through parallel traversal of each graph, which introduces its own algorithmic complications. Finally, the broader utility of GRG still depends on adoption by the genomics tool ecosystem. *snputils* currently reads GRG as an input format but does not compute on it ^22^, which is a good first step for adoption, and integration with other frameworks could accelerate community uptake. While GRG-based PCA already uses the least RAM compared to other PCA methods, traversing the graph directly from disk would be an even lower-RAM approach (at the cost of disk access during traversal) which could enable scaling to datasets an order of magnitude larger than current WGS datasets.

Taken together, GRG v2 and *grapp* demonstrate that moving from tabular to GRG-based representations can deliver substantial gains in speed, memory, and cost, while leveraging the rich Python scientific computing ecosystem. The bigger opportunity is methodological: once computational cost is no longer a bottleneck, analytical choices can be made based on their statistical justifications rather than computational feasibility. The LOCO PCA approach is one small example of a method that only becomes appealing once it is cheap to run, and we expect many more such methods to emerge in the GRG ecosystem as biobank WGS data becomes the default starting point for statistical genetics.

## Resource Availability

### Lead Contact

Requests for further information and resources should be directed to and will be fulfilled by the lead contact, Xinzhu Wei.

### Materials Availability

This study did not generate any materials.

### Data and Code Availability

This study did not generate any genetic data. The two software packages presented in this study are available at https://github.com/aprilweilab/grgl and https://github.com/aprilweilab/grapp.

## Acknowledgments

This work is partially supported by NIH R35-GM150579 and NSF IIS-2435801. This research is conducted under the UK Biobank application 97908.

## Author Contributions

Conceptualization, D.D. and X.W.; Methodology, D.D., C.A., and X.W.; Software, D.D. and C.A.; Formal analysis, D.D., C.A., and Z.P.; Investigation, D.D., C.A., and Z.P.; Visualization, D.D. with the input of all authors; Writing – original draft, D.D. and X.W.; Writing – revision, D.D. and X.W.; Writing – review & editing, all authors; Supervision, X.W.; Project administration, X.W.; Funding acquisition, X.W.

## Supplementary Information

Document S1: Supplementary methods, Tables S1-S8, and Figures S1-S14.

## Methods

### Experimental Model and Study Participant Details

We used a pre-existing UK Biobank dataset for this work. For the GWAS-with-covariates comparisons (including PCA for controlling population stratification), we used a smaller dataset of unrelated white British individuals, and removed all ultra-rare variants (allele count < 20), resulting in 338,117 individuals with 89,988,512 total variants. White British individuals were identified based on self-reported ancestry (UK biobank field 22006), after excluding individuals with excessive missing genotypes, heterozygosity or discordance between self-reported and genetically inferred sex. Pairwise relatedness was estimated using *PLINK2* (*--make-king-table*) based on high-quality array SNPs filtered for missingness <5%, minor allele frequency >1% and Hardy-Weinberg equilibrium (p-value > 10^-8^). We first identified all the related individuals with kinship threshold >=0.04419417 (corresponding to <= 3rd-degree relationships) and used them to construct a maximally independent/unrelated set by using *networkx.maximum_independent_set* in python. We further took the set union of it with individuals who have more than 10 relatives. All individuals within the set union were removed to form the unrelated white British. Since this dataset was phased by Beagle, which also imputed missing genotypes, we did not exclude any variants or samples due to missingness. The variants for the GWAS and PCA comparisons were not filtered according to Hardy-Weinberg equilibrium thresholds.

## Method Details

### Improved Graph Construction

GRG construction has two required steps (*Build* and *Merge*) and two optional steps (*Reduce* and *Simplify*). *Build* is the step that creates a graph, the remaining steps operate on existing graphs. The algorithm proceeds by splitting the genome into large chunks. Within each large chunk, *Build* is called multiple times, once for each subchunk to create a multi-tree, and then the trees within the large chunk are combined into a single graph with *Merge. Reduce* is (optionally) run on the merged graph, then *Simplify* occurs during writing the graph to disk. Finally, the graphs for all chunks are combined with *Merge* and *Simplify*. In GRG v2 we have improved *Build* (see Results section above) and added the *Reduce* step.

When graphs get merged, many nodes can end up sharing the same subset of children. The new *Reduce* step shrinks the graph by adding hierarchy in these scenarios. Given a node *n* with *k* sibling nodes *s*_*i*_, *0* ≤ *i* ≤ *k*, it chooses the sibling *s*_*i*_ with the largest number of children that are shared with *n*. If *n* and *s*_*i*_ share a sufficient number of children then creating a new parent node for these shared children will reduce the total number of edges in the graph, at the cost of adding one node. Creating hierarchy optimally between *n* and its siblings is very computationally expensive; instead we perform this pairwise algorithm for every node in the graph, optionally *reducing* the graph more than once.

GRG v1 required user specified inputs such as the number of trees to build across the genome. Determining a reasonable number of trees could be challenging, and using equal sized regions by number of variants or by length is a poor proxy for the haplotype variation that exists across the genome. We now track the number of unique haplotype segments observed across all samples, over a given region. Using linear regression, we can predict the (logarithmic) relationship between the number of samples and the expected number of unique haplotype segments per tree, to get a more “optimal” region length that each tree should span, which reduces overall graph size.

GRGs can now be constructed with different “compression” levels: 1 through 9. Higher numbers run *Reduce* for more iterations, and lower numbers increase the number of trees spanning a region, which can speed up construction while producing less optimal graphs. The file format and data structure are agnostic to the compression level, it only affects the composition of nodes and edges in the graph.

The nodes and edges of the GRG are stored in Compressed Sparse Row (CSR) format. All the edges are in a single array, and each node indexes into that array to determine which sub-array represents the edges for that node. We only store the downward (parent to child) edges in the GRG file, and each edge is represented by the *NodeID* of the child. By storing the edges in ascending order of *NodeID* within each sub-array, they can be efficiently integer encoded using *libvbyte*^*35*^ on the differences between ordered *NodeID*s. In large datasets, this reduces the graph size by about half, for both the on-disk and in-RAM graph.

### Graph-based matrix multiplication

Let *N*_*TOTAL*_ be the number of nodes in the GRG *G* (representing haploid genotype matrix *X*), where the nodes are indexed 0…(*N*_*TOTAL*_-1). Let the node ordering 0…(*N*_*TOTAL*_-1) reflect a bottom-up topological order of nodes, such that visiting all nodes < *k* ensures that all children of node *k* have been visited. Conversely, visiting all nodes > *k* ensures that all parents of node *k* have been visited, so (*N*_*TOTAL*_-1)…0 is a top-down topological order. Let *n*_*v*_(*i*) be a numerical value (initialized to 0) for each node *i*. Given the variants of *G* (i.e., the columns of *X*) numbered 0…(*M*-1), let *M*_*G*_ be the ordered list of node indexes corresponding to variants 0…(*M*-1). Recall that the *N* samples are mapped to nodes 0…(*N*-1).

Let *N*_*i*_ be the number of children for the *i*^th^ node, and *C*_*ij*_ is the node number for the *j*^th^ child of the *i*^th^ node. Then the dot-product *y* = *vX* can be computed by first copying the input vector to the sample nodes via *n*_*v*_(*i*) = *v*_*i*_ for *0* ≤ *i* < *N*, then applying the recurrence 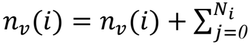 to nodes *i* in order 0…(*N*_*TOTAL*_-1), and finally *y* = *n*_*v*_(*i*), *i* ∈ *M*_*G*_, extracting the per-variant values (Figure 1c). The dot-product *y* = *vX*^*T*^ is computed by copying *v* to the variant node values, and extracting *y* from the sample node values, while the recurrence becomes *n*_*v*_(*C*_*ij*_) = *n*_*v*_(*C*_*ij*_) + *n*_*v*_(*i*) for *0* ≤ *j* < *N*_*i*_ for each node *i* in order (*N*_*TOTAL*_-1)…0.

Extending this to the matrix-matrix product *Y* = *AX* (where *A* has dimensions *K* × *N*) is trivial: *n*_*v*_(*i*) becomes a column vector of length *K* instead of a scalar, and all operations above become element-wise operations on these column vectors (Figure 1d).

The GRG library *pygrgl* implements these operations in the *matmul()* function.

### Linear Operators

The *scipy.sparse.linalg.LinearOperator* interface ignores matrix representation (e.g., dense, sparse, or GRG) by abstracting operations to matrix-matrix or matrix-vector multiplications (the *_matmat*(), *_rmatmat*(), *_matvec*(), and *_rmatvec*() functions). This is convenient for performing matrix multiplication in numpy, such as *X @ A* (where @ is matrix multiplication in Python), but the real benefit is in the following transformations.

#### Diploid genotype matrix

Let *G* be a GRG that supports haplotype-based matrix multiplication as *G*^*UP*^ (*N* × *M* matrix) and *G*^*DOWN*^ (*M* × *N* matrix). Then the vector-matrix product *y* = *vX*, where ***X*** is the *diploid* genotype matrix, can be computed by *y* = *uG*^*UP*^ where *u* is length *2 dim*(*v*) = *2N* and each diploid value *v*_*i*_ is copied to *u*_*2i*_ and *u*_*2i*+*1*_. In both cases *dim*(*y*) is *M* (number of variants). The transpose product *y* = *vX*^*T*^ is computed by *w* = *vG*^*DOWN*^ where w is length *2 dim*(*y*) and *y*_*i*_ = *w*_*2i*_ + *w*_*2i*+*1*_.

#### Standardized genotype matrix

Let ***X*** be the standardized genotype matrix, then *vX*^*T*^ = (*Xv*^*T*^)^*T*^ is computed from *G*^*DOWN*^ as described previously ^21^. *vX* = (*X*^*T*^*v*^*T*^)^*T*^ is similarly computed from *G*^*UP*^, see Supplemental Section 2.3.1 for details. We standardize using *pf*_*i*_ for the per-variant mean, where *p* is the ploidy of the genotype matrix and *f*_*i*_ is the allele frequency for variant i. Scaling can be done using multiple methods: the *sqrt* of the binomial variance *pf*_*i*_(*1* − *f*_*i*_) is the default, but the *sqrt* of the sample variance (Supplement section 2.4) can also be used. For a given variance *v*_*i*_ the alpha model ^36^ can be used, which scales multiplicatively according to *sqrt*(*v*_*i*_^*α*^) (and *α* = −*1* is equivalent to the default scaling).

#### Sample and variant covariance matrices

Once the other operators are defined, the product *vXX*^*T*^ can be computed by (*v @ op*(*X*)) *@ op*(*X*^*T*^), where *op()* is a GRG-based LinearOperator and *@* is the numpy matrix multiplication symbol. *vX*^*T*^*X* is similarly computed by (*v @ op*(*X*^*T*^)) *@ op*(*X*).

#### Multi-GRG operators

There is a wrapper around each of the above LinearOperators that takes a list of GRGs instead of a single GRG. This treats each matrix multiplication as if all of the GRGs were a single GRG, e.g. a multiplication against all autosomes combined can be done with each autosome in a separate GRG file. Each GRG can be processed in its own thread.

### GWAS with Covariates

Under a linear mixed model let *y* = *Xβ*_*G*_ + *Cβ*_*C*_ + *1β*_*0*_ + *ϵ*, where *X* is the genotype matrix, *y* is a vector of phenotypes, and *C* is a matrix of covariates. Then if we move the intercept into the covariates *C*_*I*_ = *Cβ*_*C*_ + *1β*_*0*_, we can use QR-decomposition of *C*_*I*_ to produce an equation *y*_*adj*_ = *X*_*adj*_*β*_*G*_ + *ϵ*_*adj*_ where the covariates have been factored out of both *X* and *y* (see Supplement for details). We are using simple linear regression on *X* thus letting *x* refer to a single column of *X_adj_* we get 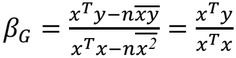 because *X*_*adj*_ has had the intercept (for both *X* and *C*) removed by adjusting for *C*_*I*_. Then we need only the diagonals of two matrix products: 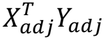 and 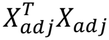. We then use one GRG multiplication for 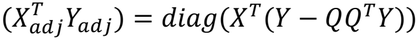 and two GRG multiplications for 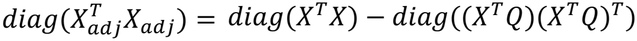. Standard errors and p-values are computed from *X*_*adj*_ in the same way as they would be from *X*.

### Principal Component Analysis

PCA of the genotype matrix X can be performed via singular value decomposition (SVD) of the standardized *X*, where *X* = *UΣV*^*T*^, and the first *k* principal component (PC) scores are the first *k* columns of *U*. PCA can also be formulated in terms of the eigen decomposition of the square matrix *XX*^*T*^ = *PDP*^*−1*^ where *P* are the eigenvectors, and *D* is the diagonal matrix containing the eigenvalues, and the first *k* columns of *P* are the PC scores. Alternatively, we can eigen-decompose *X*^*T*^*X* = *QΛQ*^*−1*^ and compute the PC scores as *Q*_*k*_*XS* where S is a diagonal matrix such that 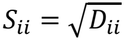. By default, *grapp* performs the eigen decomposition of *XX*^*T*^ because *N* is typically much smaller than *M* for WGS data. The user can optionally request to obtain the eigenvectors and eigenvalues, in which case *X*^*T*^*X* is decomposed. We use *scipy.sparse.linalg.eigsh()* for eigen-decomposition, which uses the implicitly-restarted lanczos method^37^.

We also implemented ProPCA ^12^ in *grapp*, which formulates the PCA problem probabilistically, and uses an expectation-maximization algorithm to find a low-dimensional approximation to *X*. SVD on this approximation produces estimates of the principal components of *X*.

PCA makes use of the multi-GRG operators to perform whole-genome multiplications, enabling thread-based parallelism and transparently handling an arbitrary number of chromosomes/graphs.

### Missing Data

While the simulated and UK Biobank data used in this paper contained no missing genotypes, the methods presented here *do* support missing phenotypes and genotypes. Missing phenotypes are supported by the GWAS operations (the matrix multiplication masks the corresponding individuals in the GRG).

Missing genotypes are supported by the LinearOperators, which optionally impute missing alleles using the allele frequency. The missing data is stored at the haplotype level in GRG, even when the missingness occurs at the individual (e.g., diploid) level.

In GRG, all variants are stored as mutations: a position (site) plus the reference allele and a single alternate allele. Each mutation has associated with it a node (the “mutation node”). When a dataset has missing genotypes, each mutation can also have another node associated with it: the “missingness node”. Samples (haplotypes) reachable from the missingness node have missing data for the site associated with that mutation. For multi-allelic sites, if any of the mutations have a missingness node then all of them will share the same missingness node. *X* is the matrix containing 0s for missing data; whereas, *X*_*M*_ is the matrix containing 1s for missing data and 0s everywhere else. Neither *X* nor *X*_*M*_ are ever formed; they are implicitly represented by the GRG. The genotype matrix *X’* should contain 0s for not having a variant, 1s for having a variant, and imputed values for having a missing variant. Assuming missingness is imputed to a fixed value per variant, we have the relationship 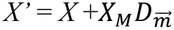, where 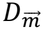 is a diagonal matrix representing imputed values. We now have 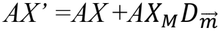 and 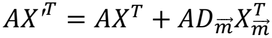. Here, *AX* and *AX*^*T*^ can be computed via naive GRG matrix multiplication. To compute 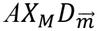 GRG matrix multiplication (“*matmul()*”) can produce a matrix *AX*_*M*_ which contains values for the missing data *only*. In the other direction, *matmul()* for *AX*^’*T*^ can consume a matrix 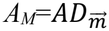 and the resulting computation actually becomes 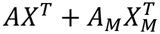. The linear operators make use of this *matmul()* mechanism by taking an optional input vector 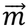 and then (implicitly) forming matrix 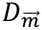 with 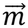 on the diagonal. With these mechanisms, we mean-impute missing data for GWAS, PCA, and other calculations, by setting 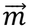 to the allele frequencies (Supplemental Section 2.4).

All of the LinearOperators in *grapp* support mutation filters and sample filters, which allows client code to ignore sets of variants and samples, and changes the dimensions of the implicit matrix that the LinearOperator represents. For example, an *N* × *M* genotype matrix represented by GRG *G* will become a *N′* × *M′* matrix via Op(*G*) when Op() has mutation filters that keep *M*’ variants and *N*’ individuals. Missing phenotypes during GWAS are treated like they do not exist in *Y* or *X*, by filtering them out with the LinearOperator and removing them from the phenotype vector *Y*. Since we are ignoring some individuals, the coalescence counts in the GRG no longer exactly represent the (current) sample variance. However the sample variance of this larger dataset (without samples filtered out) should still provide an accurate variance estimate. Alternatively, the binomial estimation of the variance (Supplement section 2.4) can be used.

### Filtering Data

GRG can optionally store population labels for sample nodes, and if present they can be used for filtering (e.g, to keep only samples from *pop1*). Samples can also be filtered by individual identifier or haplotype index. A GRG has a ploidy *p* associated with it, and filtering by haplotype index will change the GRG ploidy to 1 if the user filters partially from individuals (given ploidy *p*, each *p* consecutive haplotypes belong to the same individual).

*grapp* can filter variants based on the Hardy-Weinberg equilibrium exact test ^38^, variant type (SNP, MNP, INDEL, etc.), frequency, count, position, or number of alleles (e.g., keeping only bi-allelic sites). From the Python API, an arbitrary variant or sample-based filter can be used.

### UK Biobank analyses

In the UK Biobank cloud platform, the GRG PCA used node type *mem3_ssd1_v2_x32*. For constructing GRGs, the node types *mem1_ssd1_v2_x72* (for the 200k individuals dataset) and *mem2_ssd1_v2_x64* (for the 500k individuals dataset) were used. Supplementary Table S4 lists the command line options that were used when constructing GRGs.

We compared the use of the top 20 PCs from three different methods: GRG-based PCA on all 22 autosomes for this dataset (“ALL”), GRG-based PCA on all 21 autosomes excluding the target chromosome (“LOCO”), and the PCA results from UK Biobank field 22009 that were generated after LD pruning (147,604 total variants used). We examined chromosomes 3, 5, 6, 7, 8, 9, 10, for a mix of chromosomes that showed LD effect on the top 20 PCs (chromosomes 5, 6, 7, 8, 10) and ones that did not (chromosomes 3, 9).

We used three subsets of the UK Biobank data in our results. GRGs were constructed for the entire dataset of 490,541 individuals and 706,556,181 variants. PCA was run on all individuals after filtering the GRGs to alleles with count of 20 or greater (490,541 individuals and 137,116,837 variants). Finally, GWAS comparisons made use of only white British individuals, as described above (338,117 individuals and 89,988,512 variants).

### Tests using simulated data

Our PCA testing against simulated data used *stdpopsim* with model OutOfAfrica_4J17 and the *msprime* engine. Human recombination rate maps HapMapII_GRCh38 ^39^ were used, along with gene conversion parameters *gene_conversion_fraction* = 0.02 and *gene_conversion_length* = 300. An equal number of samples were taken from each of the four populations. Simulated data for the GWAS *β* and p-value comparisons used the same model and parameters, but only sampled a single population instead of all four. Phenotypes for the GWAS *β* and p-value comparisons were generated using *grg_pheno_sim* ^21^ with *h*^*2*^ = *0.33*, using all the variants in the dataset as causal variants (sampling their effects from a normal distribution). All GRGs were constructed from tabular data (IGD or .*vcf.gz*) exported from the simulation, not converted from the simulation ARG, as this provides a more fair comparison with other file formats.

All experiments with simulated data were performed on a server with two AMD EPYC 7532 32-Core Processors (64 regular cores, 128 hyperthreaded cores, AVX2) and 1TB of RAM.

### Threading/Parallelism

Currently, GRG-based operations are single-threaded when operating on a single graph. GRG construction is highly parallel because it decomposes each chromosome into many chunks that result in many graphs, which are then merged. Once GRGs are constructed, you can split them again (along the chromosome) to parallelize operations, but the experiments described here operate on one GRG per autosome, meaning that up to 22 threads may be used.

