## Supplement for "General, orders-of-magnitude faster whole-genome analysis with genotype representation graphs"

**5 Tables** **15**

**6 Figures** **23**

### 1 GRG Construction

GRG construction has two required steps (*Build* and *Merge*) and two optional steps (*Reduce* and *Simplify*). *Build* is the only step that creates a graph, the remaining steps modify existing graph(s). The algorithm proceeds by splitting the genome into large chunks, for the purposes of parallelization - typically each chromosome is split into at least 100 parts. The user can choose a specific number of parts, or let the algorithm choose based on default settings and the number of threads to be used. Within each large chunk, *Build* is called multiple times, once for each tree, and then the trees are combined into a single graph with *Merge*. *Reduce* is (optionally) run on the merged graph, then *Simplify* occurs during writing the graph to disk. Finally, the graphs for all parts are combined with *Merge* and then *Simplify* again runs during writing to disk.

#### 1.1 Build

The *Build* algorithm performs neighbor joining of all haplotypes, and then all parent nodes of those haplotypes, and so on, in a way that can produce unbalanced trees. Hamming distance between haplotypes (mutations present or not) is used as a distance measure, and the nearest neighbor is found by using a BK-Tree index [2].

#### 1.2 Haplotype segment representation

One challenge is fitting a large number of haplotypes in memory during each Build call. When the dataset contains many samples, loading all haplotypes can be prohibitive. We use a very simple compression technique that loads variants in chunks of 254 that we call "haplotype segments." These haplotype segments are stored as bit vectors, but we do not store one bit vector per sample (haplotype), we store only the uniquely encountered haplotype segments. For short regions (e.g., 254 consecutive variants) and large sample sizes there will be many haplotypes that are identical, and we map them all to the same haplotype segment in RAM and give it an identifier. The actual samples are then a vector of $K$ haplotype segment identifiers, where $K = L/254$ for a tree that will span $L$ variants. This representation, illustrated in Supplementary Figure S1, saves approximately $8\times$ the RAM over the bitvector representation of the full haplotypes, and also improves the performance our nearest neighbors calculations because (a) many comparisons will involve haplotype segments of the same ID, which we know are identical and (b) we can cache distance calculation results for (some) pairs of haplotype segment IDs.

#### 1.3 Limited multi-tree and mutation mapping

Previously [6], *Build* consisted of two steps: *BuildShape* and *MapMutations*. They were done separately because *BuildShape* relied on an approximate haplotype encoding to build the trees, and could not properly map the mutations during tree building. Now that we have an exact haplotype representation that is just as RAM-friendly as the previous representation, we can map the mutations during tree building and no longer call *MapMutations*.

While it is helpful to think of *Build* as creating trees, it actually creates a limited multi-tree: each node in the tree can have up to three parents, only one of which continues on upwards in the tree (the "primary parent"), while the other two serve as placeholders for mutations that cannot be represented by the primary parent. Each internal node has an "internal haplotype" associated with it, which is the intersection of the haplotypes of its children for the primary parent, and the set subtraction for results are stored in the extra parent(s) (if present). Doing so maintains a lossless mapping between mutations and samples (haplotypes). After tree construction completes, the root nodes are traversed and (newly created) mutation nodes are connected to any root node containing that mutation in its internal haplotype.

#### 1.4 Variable-sized integer encoded edge lists

The nodes and edges of the GRG are stored in a Compressed Sparse Row (CSR) format. In this format, all the edges are in a single array, and each node indexes into that array to determine which sub-array represents the edges for that node. We only store the downward (parent to child) edges in the GRG file, and many algorithms (including matrix multiplication) do not require "up" edges (child to parent). When up edges are required, they are reconstructed in RAM during the load of the down edges. Each edge is represented by the *NodeID* of the target (child). We store the edges in ascending order of *NodeID*, within each sub-array of edges. These sub-arrays can then be efficiently integer encoded in a variable-length way, using libvbyte [9] on the differences between subsequent *NodeIDs*. For example, if a node $n$ has edges to other nodes (represented by their numeric *NodeIDs*) $100, 150, 9000, 9011, 9990$ then we will store the list as $100, 50, 8850, 11, 79$. Four of these numbers can be stored using only 8 bits, and one requires 16 bits. In large datasets, we find this reduces the overall graph size by about half – since the on-disk and in-RAM representation of the graph nodes and edges is identical, this saves RAM as well as disk space.

#### 1.5 Tree size detection

The original GRG construction algorithm required users to specify the number of trees to build across the genome, and all trees were of equal size (number of variants). It was difficult to determine how many trees were needed, and having them be equal in size did not reflect the variation in diversity across the genome. With the haplotype segment representation of haplotypes, we now have a simple measure of diversity that has been so far observed: the number of unique haplotype segments will be higher for more diverse regions. Given a target number of unique haplotype segments, the tree building algorithm moves from left-to-right along the genome until that target is reached, and then it proceeds to build a tree. It then continues on from there to build another tree, until the whole region is covered by trees. The expected number of unique haplotypes should be a function that is logarithmic in the number of samples (haplotypes). We collected GRG size information on datasets of different sample sizes, and varying the target unique haplotype segment count ($T_h$). We then chose the the $T_h$ for each dataset that generated the smallest graph, and performed linear regression $T_h = b * log(N)$, where $N$ is the number of samples. This function can be used to predict the target $T_h$ for future, unseen datasets.

#### 1.6 *Merge*

Two *Merge* algorithms are supported. The original algorithm, based on hashing the samples reachable from a node, is shown below. This algorithm can be expensive on extremely large datasets, because it has to propagate the sample lists during a graph traversal. An alternative algorithm, which is the default now, uses a one-way hash of the child NodeIDs instead of the samples. Since the algorithm is performed in topological order, any children that are shared nodes will have already been mapped to the other GRG's node IDs, thus truly identical subtrees will be mapped to each other in this merge algorithm. Subtrees that differ in topology, but are identical in the samples covered, will not be mapped to each other - this can be good for retaining hierarchy in the graph, but will result in slightly larger graphs overall.

**Algorithm 1** Merge two GRGs

```
Merge(grg1, grg2):
  hashToNode ← ∅
  for all node ∈ depthFirstOrder(grg1) do
    hashToNode ← hashToNode ∪ (node, oneWayHash(getCoveredSamples(node)))
  end for
  grg2ToGrg1Map ← ∅
  for all node ∈ depthFirstOrder(grg2) do
    hashValue ← oneWayHash(getCoveredSamples(node))
    if hashValue ∈ hashToNode then
      nodeInGrg1 ← hashToNode[hashValue]
      nodeInGrg1.mutations ← nodeInGrg1.mutations ∪ node.mutations
    else
      nodeInGrg1 ← makeNode(grg1)
      nodeInGrg1.mutations ← node.mutations
      for all child ∈ node.children do
        addEdge(grg1, nodeInGrg1, grg2ToGrg1Map[child])
      end for
    end if
    grg2ToGrg1Map ← grg2ToGrg1Map ∪ {(node, nodeInGrg1)}
  end for
```

The only difference between these two algorithms is whether the one-way hash is computed from the list of covered samples ($getCovertedSamples()$) or the list of immediate children ($getChildren()$), which have been already replaced with equivalent nodes from $grg1$.

##### 1.7 *Simplify*

A GRG can be simplified by observing the following patterns of nodes and edges that do not contribute to the compression of the graph, but simply add extra hierarchy:

1. Any node $n$ with a single incoming edge from its parent $p$. $n$ can be removed, and all of its outgoing edges can become outgoing edges of $p$ directly. This shrinks the graph by one edge and one node.

2. Any node $n$ with a single outgoing edge. This is the same as the previous pattern.

3. Any node $n$ that has two incoming edges (from parents $p_1$, $p_2$) and two outgoing edges (to children $c_1$, $c_2$). These four edges can be changed to be the following four edges: $p_1 \rightarrow c_1$, $p_1 \rightarrow c_2$, $p_2 \rightarrow c_1$, $p_2 \rightarrow c_2$, and $n$ can be removed, shrinking the graph by one node.

We will only remove a node $n$ that contains no mutations mapped to it.

##### 1.8 *Reduce*

When graphs get merged, it can result in many nodes sharing the same children. We can shrink the graph by adding hierarchy in these scenarios. Given a node $n$, find all of its sibling nodes $s_1$, $s_2$, ..., $s_k$. At each sibling, count the number of children that are shared with $n$, and choose the sibling $s_i$ with the largest count. If $n$ and $s_i$ share a sufficient number of children then creating a new node $h$ to be the parent of these shared children will reduce the number of edges in the graph, at the cost of adding one node. In general, finding the optimal way to create hierarchy between $n$ and its siblings is very computationally expensive, so we just perform this pairwise algorithm for every node in the graph and iterate over the graph multiple times until the graph is no longer shrinking.

#### 2 GRG matrix multiplication

##### 2.1 Dot product

Given a GRG $\mathbf{X}$ with $t$ nodes (representing a $n \times m$ genotype matrix), assume that we can store a numerical value at each node, and that $v_i$ is the value of the $i^{th}$ node. Given a vector $u$ and a GRG $\mathbf{X}$, the matrix-vector product $y = u \times \mathbf{X}$ is computed by by first setting the value of each sample node $j$ ($1 \leq j \leq n$) with the corresponding $u$ value. That is, set $v_j = u_j$. Then visit each of the $t$ nodes in bottom-up topological order (so that all children are visited before each parent node), and set $v_i = \sum_{c \in children(i)} v_c$. I.e., the parent value is equal to the sum of the values of its children. Let $g(i)$ be the node number associated with the $i^{th}$ mutation. Then for each mutation (variant) in the GRG numbered $1...m$ (in order of ascending genetic position), copy the mutation's value into the result vector, i.e. $y_i = v_{g(i)}$ for $1 \leq i \leq m$.

##### 2.2 Matrix-matrix multiplication

Given a $k \times n$ matrix $\mathbf{A}$, the matrix multiplication $\mathbf{Y} = \mathbf{A} \times \mathbf{X}$ is just the concatenation of the $k$ matrix-vector product $\mathbf{A}_i \times X$ for each row $i$ ($1 \leq i \leq n$). To perform this on a GRG, assume we have a vector of values $v_i$ at each graph node, instead of a scalar value. The graph traversal is identical to the dot product, except that now we initialize the *vector* $v_j = A_{:,j}$ (the $j^{th}$ column initializes the $j^{th}$ sample node's value vector). Similarly, each parent value adds the vectors of its children into a vector result. Figure S6 illustrates the matrix multiplication process for $\mathbf{Y} = \mathbf{A} \times \mathbf{X}$. Note that $\mathbf{Y} = \mathbf{A} \times \mathbf{X}^T$ is just performed by traversing in the opposite direction (from mutations to samples).

Matrix multiplication can be performed on haploid samples or individuals with ploidy $p$. In the latter case, an individual's value is found by summing the values associated with its sample (haplotype) nodes. The matrix multiplication method (*pygrgl.matmul*) provides a flag to toggle this behavior (*by_individual=True*).

The matrix $X$ represented by the GRG is a matrix with only the values 0, 1. A special case of matrix multiplication is when the input matrix $\mathbf{A}$ also only takes on values 0, 1, and there is only one non-zero value per row. Such a matrix multiplication is equivalent to asking for all the samples associated with a single mutation, or all the mutations associated with a single sample. Such a multiplication can recover the *explicit* form of the genotype matrix from the GRG, and in fact we use this to export a GRG to the IGD [5] format. For this reason, GRGL supports bit-packing the node value vectors $v_j$, and efficient bit-wise operations to perform the per-node "addition" step.

#### 2.3 Linear operators

GRG's matrix multiplication enables iterative linear algebra methods on huge genetic datasets. *grapp* implements multiple operators that abstract away the details of multiplications against the genotype matrix, and comply with the `scipy.sparse.linalg.LinearOperator` interface for integration with existing *scipy* APIs for eigen decomposition, singular value decomposition (SVD), and a variety of methods for solving systems of linear equations (e.g., conjugate gradient). `LinearOperator`s are a way to implicitly represent a matrix, by supporting matrix-matrix and matrix-vector multiplication against a possibly transformed version of the target matrix.

The simplest operator is the `SciPyXOperator`, which is equivalent to matrix multiplication against the GRG. It, and all other `LinearOperator`s, can support multiplication against the $0, 1$ valued matrix (haplotypes) or the $0, 1, 2$ valued matrix for diploid individuals (in fact it can support ploidies larger than 2).

The `SciPyXTXOperator` allows you to perform multiplications against the $m \times m$ matrix $\mathbf{X}^T\mathbf{X}$. The `SciPyXXTOperator` similarly allows multiplication against the $n \times n$ matrix $\mathbf{XX^T}$.

##### 2.3.1 Standardized operators

The standardized genotype matrix is formed by subtracting the mean from each column (mutation) and dividing by the standard deviation. Since traditional (small dataset) calculations explicitly represent the genotype matrix, they can transform it into the standardized matrix directly. The storage (and computation) cost for both representations is roughly the same. The GRG stores the genotype matrix in a sub-sparse matrix encoding, and this is only possible because of the $0, 1$ valued nature of the matrix, therefore standardizing the GRG cannot be done explicitly, it has to be done *implicitly*. For this reason, we provide LinearOperators that multiply against the standardized genotype matrix.

We illustrate the standardized operators by deriving the product

$$\mathcal{X}^T \mathcal{X},$$

where

$$\mathcal{X} = (X - U)\Sigma$$

is the standardized diploid genotype matrix. $X \in \mathbb{R}^{n \times m}$ is the genotype matrix, $U \in \mathbb{R}^{n \times m}$ has the $i$th column equal to $2f_i$, where $f_i$ is the allele-frequency of mutation $i$ and $\Sigma \in \mathbb{R}^{m \times m}$ is diagonal with

$$\Sigma_{ii} = \frac{1}{\sigma_i}, \qquad \sigma_i = \sqrt{2f_i(1 - f_i)}.$$

For a `scipy.sparse.linalg.LinearOperator`, all we need is a way to compute

$$y \;=\; (\mathcal{X}^T \mathcal{X})v \;=\; \mathcal{X}^T(\mathcal{X}v)$$

for any given vector $v \in \mathbb{R}^m$. We do this in two steps.

**Step 1: Compute $\mathcal{X}v$**

$$d = \mathcal{X}v = (X - U)\Sigma\, v = X(\Sigma v) - U(\Sigma v).$$

We observe that

$$w = \Sigma\, v, \quad w_i = \frac{v_i}{\sigma_i}.$$

Then for each sample index $j = 1, \ldots, n$,

$$d_j = (Xw)_j - \sum_{i=1}^{m}(2f_i)w_i = (Xw)_j - \sum_{i=1}^{m} 2f_i \frac{v_i}{\sigma_i}.$$

**Step 2: Compute $\mathcal{X}^T d$**

$$y = \mathcal{X}^T d = \big((X - U)\Sigma\big)^T d = \Sigma\,(X^T d) - \Sigma\,(U^T d).$$

We observe that

$$U^T d = \begin{pmatrix} 2f_1 \sum_{j=1}^{n} d_j \\ 2f_2 \sum_{j=1}^{n} d_j \\ \vdots \\ 2f_m \sum_{j=1}^{n} d_j \end{pmatrix}$$

Let $s = \sum_{j=1}^{n} d_j$, then we get

$$(U^T d)_i = 2f_i\, s, \quad (\Sigma\, U^T d)_i = \frac{2f_i}{\sigma_i}\, s.$$

Thus we can get each element of the desired vector,

$$y_i = \frac{(X^T d)_i}{\sigma_i} - \frac{2f_i s}{\sigma_i}$$

The `SciPyStdXTXOperator` computes the above product (against the correlation matrix) in this way. `SciPyStdXOperator` computes *Step 1* (the product $\mathcal{X}v$) or *Step 2* (the product $\mathcal{X}^T d$) depending on which one the user chooses. Similarly, there is a `SciPyStdXXTOperator` operator for performing multiplications against the genetic relatedness matrix (GRM).

##### 2.3.2 Multi-GRG operators

Often calculations like PCA or GWAS need to be computed over all of the autosomes simultaneously. GRGL provides exended versions of the `LinearOperator`s that take lists of GRGs (e.g., one per chromosome) and can perform a single multiplication in parallel over the graphs.

#### 2.4 GWAS with covariates

At a single mutation, let

$$\mathbf{x} = (x_1, \dots, x_n)^T \in \mathbb{R}^n \quad \text{and} \quad \mathbf{y} = (y_1, \dots, y_n)^T \in \mathbb{R}^n$$

be the vectors of diploid genotypes and phenotypes, respectively, so that each $x_i \in \{0, 1, 2\}$. We compute simple linear regression GWAS with the following equation

$$\beta \;=\; \frac{\sum_{i=1}^{n}(x_i - \bar{x})(y_i - \bar{y})}{\sum_{i=1}^{n}(x_i - \bar{x})^2} \;=\; \frac{\mathbf{x}^T\mathbf{y} \;-\; n\,\bar{x}\,\bar{y}}{\mathbf{x}^T\mathbf{x} \;-\; n\,\bar{x}^2}.$$

Let

$$\mathbf{X} = \begin{pmatrix} \mathbf{x}_1^T \\ \mathbf{x}_2^T \\ \vdots \\ \mathbf{x}_n^T \end{pmatrix} \in \mathbb{R}^{n \times m},$$

where each $\mathbf{x}_i^T = (x_{i1},\, x_{i2},\, \dots,\, x_{im}) \in \mathbb{R}^{1 \times m}$ is the $i$-th row of $\mathbf{X}$, the full genotype matrix. In GRG, the two dot products of $x^T x$ and $x^T y$, are computed for each row vector $\mathbf{x_n}$ via graph traversals.

$$x^T x = \text{sample count} + 2 \times \text{homozygote count}$$

is computed by using the stored value for the number of individuals that coalesce at a given node. This information is propagated from the children nodes to the parents to yield a homozygote count as found in a traditional diploid matrix. $x^T y$ is a dot product that can be found through one graph traversal.

Now we extend this to control for covariates. Under a mixed linear model, let:

$$Y = X\beta_G + C\beta_C + \epsilon \tag{1}$$

$$X \in \mathbb{R}^{n\times m}, \quad C \in \mathbb{R}^{n\times c}, \quad Y, \epsilon \in \mathbb{R}^n, \quad \beta_G \in \mathbb{R}^m, \quad \beta_C \in \mathbb{R}^c.$$

where $X$ is the genotype matrix, $Y$ is a vector of phenotypic values, $\beta_G$ is the fixed effect that comes from genetic factors, $C$ is the covariate matrix, $\beta_C$ is the regression coefficient of the covariates, and $\epsilon$ is the error term. The more general form contains the intercept $\beta_0$, i.e. $Y = X\beta_G + C\beta_C + \beta_0\epsilon$. We assume that $\beta_0$ is included in the matrix $\beta_C$ by always including a column of 1s in $C$. We define

$$H_C = C(C^TC)^{-1}C^T \qquad P_C = I - H_C$$

such that

$$\begin{aligned} P_C C\beta_C &= (I - H_C)C\beta_C \\ &= C\beta_C - H_C C\beta_C \\ &= C\beta_C - C(C^TC)^{-1}C^TC\beta_C \\ &= C\beta_C - C\beta_C = 0 \end{aligned}$$

Multiplying the model (1) by $P_C$ gives

$$Y_{\text{adj}} := P_C Y = P_C X\,\beta_G + \underbrace{P_C C\,\beta_C}_{=0} + P_C\epsilon =: X_{\text{adj}}\,\beta_G + \epsilon'.$$

Then, we can derive $\beta_G$ as:

$$\hat{\beta}_G = (X_{adj}^T X_{adj})^{-1} X_{adj}^T Y_{adj} \tag{2}$$

We can take a QR decomposition such that:

$$C = \widetilde{Q}\,\widetilde{R}, \quad \widetilde{Q} = [\,Q_1 \;\; Q_2\,] \in \mathbb{R}^{n\times n},\; \widetilde{R} = \begin{bmatrix} R_1 \\ 0 \end{bmatrix} \in \mathbb{R}^{n\times c},$$

where $Q_1 \in \mathbb{R}^{n\times c}$ and $Q_2 \in \mathbb{R}^{n\times(n-c)}$ have orthonormal columns and $R_1 \in \mathbb{R}^{c\times c}$ is upper-triangular and invertible. Since the bottom $(n-c)$ rows of $\widetilde{R}$ are zero,

$$C = \widetilde{Q}\,\widetilde{R} = [\,Q_1\; Q_2\,]\begin{bmatrix} R_1 \\ 0 \end{bmatrix} = Q_1\,R_1.$$

Define $Q := Q_1$ and $R := R_1$ as the reduced/thin QR where

$$C = Q\,R, \quad Q \in \mathbb{R}^{n\times c},\; Q^TQ = I_c,\; R \in \mathbb{R}^{c\times c}.$$

Substitute into $H_C$:

$$H_C = QR(R^TR)^{-1}R^TQ^T = Q[RR^{-1}(R^T)^{-1}R^T]Q^T = QI_pQ^T = QQ^T.$$

Since $X_{\text{adj}} = (I - H_C)X$, we have

$$X_{\text{adj}} = X - QQ^TX, \qquad X_{\text{adj}}^T X_{\text{adj}} = X^TX - X^TQQ^TX = X^TX - (X^TQ)(X^TQ)^T.$$

The matrix $X^TQ$ has dimension $m \times c$.

To compute $X_{adj}^T Y_{adj}$ we use

$$\begin{aligned} X_{\text{adj}}^T Y_{\text{adj}} &= (PC \cdot X)^T(PC \cdot Y) \\ &= X^T PC^T PCY \\ &= X^T PCY \quad (\text{since } PC^T = PC \text{ and } PC^2 = PC) \\ &= X^T(Y - QQ^TY)) \end{aligned}$$

The above derivation is general, in that it could be used for multiple or simple linear regression, however the latter just needs $diag(X^TX) - diag((X^TQ)(X^TQ)^T)$ instead of the full matrix. For simple linear regression, the above can be done with standard matrix operations on $Y$ and $C$, plus three graph traversals over the GRG. First, we compute $diag(X^T \times X)$, using the coalescence method described above. Then, using the non-standardized `LinearOperator`, we compute $X^T \times Q$ and $Y_{adj} \times X$.

##### 2.4.1 Binomial vs. Sample Variance

The above implicitly uses the sample variance of the dataset, via $diag(X^TX)$. GRG computes $diag(X^TX)$ using per-node coalescence counts, which track the number of individuals that coalesce *exactly* at a given node. $diag(X^TX)$ can also be approximated by observing that it is a function of the per-variant variance. Observe that

$$\frac{diag(X^TX)}{n} = \frac{\sum_{i=1}^{n}(x_i - \bar{x})^2}{n} = Var[X] + E[X]^2$$

assuming that all variants follow a binomial distribution, we get

$$diag(X^TX) = n(pf_i(1 - f_i) + (pf_i)^2)$$

where $p$ is the ploidy of the genotype matrix and $f_i$ is the allele frequency of the $i^{th}$ variant. Conversely, we can compute the sample variance of $X$ via

$$Var[X] = \frac{diag(X^TX)}{n} - E[X]^2 = \frac{diag(X^TX)}{n} - (pf_i)^2$$

*grapp* includes an option to use the binomial variance for GWAS, instead of sample variance, and also has method `grapp.util.simple.variance` for computing the sample or binomial variance directly.

##### 2.4.2 Standardized Computation

For the above GWAS, each $\beta_i$ is computed independently from the others (though simultaneously, using matrix products). We provide an option to compute this using the standardized genotype matrix $(X)$. This is done using three changes from the above:

1. $Y_{adj}$ is standardized using its sample mean and standard deviation.
2. $diag(X^T \times X)$ is no longer needed. For a given mutation $j$, let $c_j = diag(X^T \times X)_j$. Then $c_j = \sum_{i=0}^{n}(x_{ij} - \hat{x_j})^2$, which can also be written as $n \times (Var[x_j] + E[x_j]^2)$, because $Var[x_j] + E[x_j]^2 = E[x_j^2] = \frac{1}{n}(\sum_{i=0}^{n}(x_{ij} - \hat{x_j})^2)$. Since $x_j$ is standardized, $Var[x_j] = 1$ and $E[x_j]^2 = 0$, thus $c_j = n$ for all $j$.
3. We simply use the standardized `LinearOperator`, to compute $X^T \times Q$ and $Y_{adj} \times X$ instead of the non-standardized.

The standardized computation uses the same number of graph traversals: we remove one traversal (for $diag(X^T \times X)$), and add a traversal for computing the allele frequencies needed for the standardized `LinearOperators`. This standardized computation assumes that $X$ and $C$ are uncorrelated.

##### 2.4.3 Missing Genotypes

To understand how GRG stores missing data, it is useful to review how it stores non-missing genotypes. All variants are stored as Mutations: a position (site) plus the allele. Each Mutation has associated with it a Node that we call the "Mutation Node". All samples reachable (following down edges) from that Mutation Node make up the sample set of that Mutation - i.e., the samples that contain that variant in their genome. All samples unreachable from that Mutation Node either contain a different Mutation at that site (if multi-allelic) or the reference allele (if no Mutation Node for that site reaches the sample). When a dataset has missing genotypes, each Mutation can also have another Node associated with it: the Missingness Node. The samples reachable from the Missingness Node have missing data for the site associated with that Mutation. For multi-allelic sites, if any of the Mutations have a Missingness Node then all of them will share the *same* Missingness Node.

To make use of missingness information, GRG matrix multiplication $Y = AX$ can also produce a matrix $M = AX_m$ which contains the computed values for the missing data only. Here $X$ is the genotype matrix containing 0 for missing data, and $X_m$ is the matrix containing 1 only in missing data locations (and 0 everywhere else). In the other direction (*down*, via graph traversal), the matrix multiplication $Y = AX^T$ can *consume* a matrix $M$ and the resulting computation actually becomes $Y = AX^T + MX_m^T$.

We want to treat each missing genotype as the mean value. For the regular (no covariates) GWAS, we just need to replace $n$ with $n_j$, which is the number of samples *without* missing data for Mutation $j$:

$$\beta \;=\; \frac{\sum_{i=1}^{n}(x_i - \bar{x})(y_i - \bar{y})}{\sum_{i=1}^{n}(x_i - \bar{x})^2} \;=\; \frac{\mathbf{x}^T\mathbf{y} \;-\; n_j\,\bar{x}\,\bar{y}}{\mathbf{x}^T\mathbf{x} \;-\; n_j\,\bar{x}^2}.$$

This is clearly the case, because we want $(x_i - \bar{x})$ to be zero if the $i$ sample has a missing value, and it will already be zero in the $X$ matrix if we ignore missingness nodes, so we can keep it zero by ignoring it in the mean adjustments $n_j\,\bar{x}\,\bar{y}$ and $n_j\,\bar{x}^2$. For the standardized genotype matrix, instead of $\mathbf{x}^T\mathbf{x}$ we use a vector of all $n_j$.

The $diag(X^TX)$ is the only calculation in the GWAS derivation that needs this special handling. All of the other graph traversals use `LinearOperators`, so now we describe how these operators properly handle missing data.

The non-standardized operators take an (optional) vector of length $m$ that has a per-Mutation missing value. Typically, the allele frequencies will be passed in for the missing values, which treats missingness as the mean value. Place the vector $m$ on the diagonal of a matrix $D_m$. Since $X$ is the genotype matrix containing 0 in all missingness cells, and $X_m$ is the matrix containing only 1 in the missingness cells, we can ignore missingness by using $X$ (all missing alleles are treated as REF) or by using $X + X_m$ (all missing alleles are treated as ALT). Instead, we mean-impute the missing alleles using a weighted sum of $X$, $X_m$: for the product $Y = AX$ we get $Y = A(X + X_mD_m) = AX + AX_mD_m$, and likewise for $Y = AX^T$ we get $Y = AX + AD_mX_m^T$.

The standardized operators behave similarly to the $diag(x^Tx)$ scenario and just need their allele frequencies (the means used during standardization) adjusted for missingness. That is, instead of $f_j = \frac{a_j}{n}$ we use $f_j = \frac{a_j}{n_j}$, where $a_j$ is the number of haplotypes containing Mutation $j$, $n$ is the total number of haplotypes, and $n_j$ is the number of haplotypes without missing data for Mutation $j$.

##### 2.4.4 Missing Phenotypes

Missing phenotypes (values in $Y$) are treated like they do not exist in $Y$ or $X$, i.e. the handling is equivalent to removing all edges related to the samples from the GRG. There are two ways to adjust the GWAS calculation, both of which are adjustments to the $diag(X^T \times X)$ term. Since we are ignoring some individuals, the coalescence counts in the GRG no longer exactly represent the (current) sample variance. One approach is to ignore this, and use the sample variance from the larger sample: the whole graph, including the individuals that are ignored in the current GWAS due to missing phenotypes. This variance estimate is likely still accurate, assuming that missing phenotypes are randomly distributed. An alternative is to use the binomial estimation of the variance (see the discussion on standardization above), which does not require the coalescence counts from the GRG.

##### 2.4.5 Standard error and t-value

We can find the the standard error primarily using values we already calculated. Let's consider a derivation for a single $\beta$ from GWAS with covariates, this would be an arbitrary entry in $\beta_G$, that has a corresponding mutation vector $x$ with a corresponding phenotype value $y$.

The residual vector can be defined as

$$r = y_{adj} - x_{adj}\beta \quad \beta = \frac{x_{adj}^T y_{adj}}{x_{adj}^T x_{adj}}$$

$$\begin{aligned}\text{SSE} = \sum_{i=1}^{n}(y_i - \bar{y})^2 = r^Tr &= (y_{adj} - x_{adj}\beta)^T(y_{adj} - x_{adj}\beta) \\ &= y_{adj}^T y_{adj} - 2\beta x_{adj}^T y_{adj} + \beta^2 x_{adj}^T x_{adj}\end{aligned}$$

$$\text{s.e.}(\beta) = \sqrt{\frac{\text{SSE}}{(n-c-1)\times x_{adj}^T x_{adj}}} \qquad \text{c here includes the intercept dimension}$$

The t-value is

$$t = \frac{\beta}{\text{s.e.}(\beta)}$$

#### 2.5 Principal component analysis

By default, our PCA implementation passes the $XX^T$ *LinearOperator* to the `scipy.sparse.linalg.eigsh` function. The $k$ eigenvectors corresponding to the largest eigenvalues are then the top $k$ (normalized) PCs. Under the hood, this uses an iterative implicitly-restarted Arnoldi method [8] for eigen decomposition. Since we are typically dealing with datasets where $M >> N$ (huge WGS datasets have hundreds of millions of SNPs and currently around a million haplotypes), it is substantially more efficient to do PCA on $XX^T$ than $X^TX$. We observe that on the UK Biobank performing PCA with $XX^T$ uses about half the RAM of $X^TX$.

Alternatively, we could use the standardized $X^TX$ genotype matrix (correlation matrix) *LinearOperator* to get the eigenvalues and vectors associated with the $k$ largest eigenvalues. The top $k$ eigenvectors are then multiplied with the standardized $X^T$ matrix to generate the principle components (i.e., projecting the per-mutation eigenvectors back over all individuals) as a $N_I \times k$ matrix. When each of the $k$ columns are divided by $\sqrt{\lambda_i}$ (where $\lambda_i$ is the PC corresponding $i$th eigenvalue, to do unit-variance normalization) the results are equivalent to just performing eigen decomposition on the $XX^T$ matrix.

It is worth noting that small numerical differences can cause the `scipy.sparse.linalg.eigs` method to flip the sign of certain eigenvectors, which was observed in our unit tests. To validate this theory we ran an experiment where $\mathcal{X}^T\mathcal{X}$ was computed directly and then `scipy.linalg.eigs` was called. To this genotype matrix we added random noise with a magnitude $1 \times 10^{-10}$ and found that the sign of some eigenvectors was flipped. This rotation around axes is present in other tools as well, and can be observed in Figure S8 where we show the top 2 PCs from a simulated out-of-Africa dataset.

##### 2.5.1 Error Analysis

To ensure the accuracy of our method we should compute PCs directly from a dense SVD of 1,000 diploid indivduals standardized genotype matrix. This is how we calculate our ground truth unit variance PCs

$$\mathcal{X} = U\Sigma V^T$$

$$\mathcal{X}^T\mathcal{X} = (V\Sigma U^T)USV^T = VS^2V$$

This is the eigendecomposition of $\mathcal{X}^T\mathcal{X}$, so

$$XV = USV^TV = US$$

Therefore, the $k^{th}$ PC can be found from the $k^{th}$ column of $U$. Comparing 10 of *grapp*'s PCs to these ten columns we get standard errors ranging from $1.36 \times 10^{-16}$ to $5.89 \times 10^{-15}$. This is slightly better than FlashPCA2 ($1.98 \times 10^{-16}$ to $1.79 \times 10^{-8}$). *plink2* isn't really possible to compare against because their output format has limited decimal precision (standard errors are therefore on the order of $10^{-8}$ to $10^{-6}$).

##### 2.5.2 ProPCA

We also implement the ProPCA algorithm [1] in *grapp*, which provides a faster algorithm (at the cost of more RAM) than the eigen-decomposition based default PCA in *grapp*.

The aim of PCA is to find $W \in \mathbb{R}^{n\times k}$ with orthonormal columns and $Z \in \mathbb{R}^{n\times k}$ to minimize

$$\|\mathcal{X} - WZ^T\|_F$$

This minimization is basically how can you best reconstruct $\mathcal{X}$ with fewer dimensions. The ProPCA algorithm includes this to check for convergence. Consider SVD

$$\mathcal{X} = U\Sigma V^T$$

We let $\hat{W} = U_k$ where $U_k$ contains the first k columns of $U$, these correspond to the largest singular values. Probabilistic PCA uses the model $y|x, \epsilon = Cx + \epsilon$ where x is from a Gaussian distribution and $\epsilon$ is noise. The MLE of $C$ spans the $k$-dimensional principal subspace, i.e. spans the same space as the top $k$ PC vectors.

The EM algorithm is used in these types of models to approximate C. Let the standardized genotype matrix $\mathcal{X} = Y$ to not confuse it

$$X = (C^T C)^{-1} C^T Y$$
$$C = Y X^T (X X^T)^{-1}$$

The ProPCA algorithm orthogonalizes the matrix $C$ to obtain the principal components in time $O(mk^2)$, via Q-R decomposition. Let

$$C = QR$$

Then

$$b = Q^T Y, \quad b \in \mathbb{R}^{k \times n}$$
$$b = U \Sigma V^T,$$
$$\text{eigenvects} = QU_k \qquad \text{this projects } k \text{ vectors from } \mathbb{R}^k \to \mathbb{R}^m$$
$$\text{eigenvals} = \frac{\sigma_i}{\# \text{ of SNP}}$$
$$\text{PCs} = V_k$$

### 3 Applications to simulated data

#### 3.1 File construction comparisons

Table S3 shows file size and construction time comparison between PGEN and GRG on simulated datasets ranging from 100,000 to 1,000,000 diploid individuals. The latest version of GRG support compression levels from 1 to 9, similar to tools like *bzip* and *gzip.* The default is level 5, and we compare that against level 1 (called "fast" in the Table S3). Default GRG is between 13× (fewest individuals in dataset) and 47× smaller than PGEN on these simulated datasets. The fast (level 1) GRG construction is roughly 10% faster than the default (for *.vcf.gz* inputs), and between 0.27× and 2.6× larger than the default GRG. The new *Reduce* stage of constructing GRGs is especially effective at compressing simulation data, and is the largest difference between the default (level 5) and fast (level 1) compression levels.

On the UK Biobank dataset (200k individuals) GRG v2 is smaller than the XSI [10] and Savvy [7] files that we constructed in DeHaas, et al., 2025[6] (Table S7), and the improvements to construction time have made the CPU-hours for creating GRG comparable to both of those formats (about 40% more than XSI), and GRG is substantially faster for computing dot-products [6]. In this experiment, we stripped the expensive meta-data out of the BCF files before creating XSI/SAV files, using `bcftools annotate --remove FORMAT <input bcf> -O b --output <output bcf>`. We used the default GRG construction arguments (which results in compression level 5), and the following command lines to construct XSI and SAV files:

- `xsqueezeit -c -f <input> -o <output>`
- `sav import --pbwt-fields GT --sparse-threshold 0.01 --phasing full`

To further explore the compression and scaling of these file formats, we created four simulated datasets, based on the same *OutOfAfrica_4J17* stdpopsim models used elsewhere. These datasets sampled an equal number of individuals from all four populations represented by this model. The datasets are:

1. 100,000 diploid individuals.
2. 100,000 diploid individuals with noise to simulate genotyping error.
3. 200,000 diploid individuals.
4. 200,000 diploid individuals with noise to simulate genotyping error.

The noise was added to the dataset on a variant-by-variant basis. Let $C_i$ be the alternate allele count at variant $i$. With probability $p = 0.005$ we flipped reference alleles to alternate alleles according to Binomial$(C_i, p)$. With the same probability $p$ we then randomly flipped alternate alleles to reference alleles.

We then tested three configuration with different command-line flags for XSI and Savvy:

- XSI v4.0.0: `xsqueezeit -c -f <input> -o <output>`
- XSI-github: `xsqueezeit -c --variant-block-length 57344 --zstd -f <input> -o <output>`
- Savvy v2.2.0: `sav import -b 65535 --pbwt-fields GT --sparse-threshold 0.001 --phasing full`

XSI's `--variant-block-length` flag is only available in the main branch of the GitHub repo (we'll refer to as "XSI-github"), at the time of this writing, and is not available in the latest tagged release (v4.0.0, we'll refer to as "XSI"). Because XSI-github failed to run on most of our BCF files and option settings, Figure S13 shows only the one successful XSI-github run, which used a large block size and built-in zstd compression.

For each file format, we also used the zstandard tool [4] to compress the file, and reported this file size separately. The noise had a rather substantial impact on the compression provided by XSI, Savvy, and GRG (Figure S13b). On noise-free data, GRG-zst and XSI-github provide the best compression (within 7MB of each other). XSI-zst is the smallest file in the noisy dataset. In both the noise-free and noisy datasets, GRG scales the best: that is, the ratio of the GRG file sizes for 200,000 and 100,000 individuals is the smallest.

#### 3.2 GWAS comparison between grapp and plink

We compared our GWAS-plus-covariates implementation against *plink2*'s `--linear` analysis. We used `omit-ref` so that *plink2* would not flip the effects to be in terms of the minor allele, so the results were directly comparable to *grapp*. In Figure S7 we plot $\beta$ and p-values from *plink2* and *grapp* against each other. You can see that a small number of $\beta$ values differ near 0 (on the y-axis), and that the p-values are highly similar, with some slight differences near the largest (least significant) ones. These differences are due to *grapp* and *plink2* treating multi-allelic sites differently: we verified that *plink2* regresses variants at multi-allelic sites jointly, whereas *grapp* regresses each minor allele separately. We verified that these differences occur at 2,525 out of 1,379,274 compared variants all of which are multi-allelic.

These tests were run on an *msprime* simulated dataset with 25,000 individuals from a single population. The phenotype was simulated with *grg_pheno_sim*. Rare variants were filtered out using MAC $< 20$ (which is also recommended by *plink2*'s documentation).

*grapp* commands:

- `grapp filter -c 20 -C 49980 stdpop.ooa4.eur25k.grg stdpop.ooa4.eur25k.c20.C49980.grg`
- `grapp assoc -p stdpop.ooa4.eur25k.c20.grg.phen -c stdpop.ooa4.eur25k.c20.grg.pcs.tsv stdpop.ooa4.eur25k.c20.C49980.grg`

*plink2* commands:

- `plink2 --mac 20 minor --make-pgen --threads 8 --vcf stdpop.ooa4.eur25k.vcf.gz --out stdpop.ooa4.eur25k`
- `plink2 --allow-no-sex --covar iid-only stdpop.ooa4.eur25k.c20.pcs.plink.cov --glm omit-ref hide-covar --out pgen.eur25k.threads1.omitref.gwas_covar --pfile stdpop.ooa4.eur25k --pheno iid-only stdpop.ooa4.eur25k.c20.phen.plink.txt --threads 1`

#### 3.3 Load time

Load times in GRG v2 are now faster or roughly the same as the time for a single matrix-vector multiplcation (Table S8). Noise in the simulated data can affects file size substantially, which is indicative of a larger graph. Since graph load time is just reading nodes and edges in a linear manner off disk, and not interpreting them, the additional graph size has minimal impact on file load time. The matrix multiplication is interpreting/using those nodes and edges, thus graph size has a more substantial impact on runtime.

##### 3.4 Compression level

On noise-free data, higher compression levels can reduce graph file size (Figure S14). On noisy data, it appears to not affect file size substantially. In both cases, the "levels" are using different heuristics for deciding how to construct the graph, so the resulting graph sizes are not necessarily monotic with respect to compression level.

#### 4 Applications to real data

##### 4.1 UK Biobank *.vcf.gz* to IGD conversion

We created GRG files from the UK Biobank WGS data of 500,000 individuals phased via Beagle. While GRGs can be constructed directly from Tabix-indexed *.vcf.gz* files, we first convert those *.vcf.gz* files to IGD format [5], a tabular format similar in simplicity to VCF but with a smaller footprint and substantially faster traversal times. IGD stores all variants (alternative allele genotype data) as bi-allelic, similar in the way that $bcftoolsnorm$ can transform VCF files. We converted all 22 autosomes from the UK Biobank dataset to IGD in the RAP cloud environment, for a total cost of 154.37GBP (4619.62 CPU hours, over 322.03 hours of elapsed time). We used the cheapest node type that provided 8 or 16 threads for the conversion, while also containing enough storage for the input (*.vcf.gz*) and output (IGD). We used SSD-based nodes, i.e. the *mem**_*ssd**_*v2*_*x8* and *mem**_*ssd**_*v2*_*x16* node types. The input *.vcf.gz* files totaled just under 3TB in size while the resulting (LZ4-compressed) IGD files were 1.18TB in size. Across all autosomes $706,556,181$ variants are stored in the IGD files.

##### 4.2 UK Biobank IGD to GRG conversion

We used the default (level 5) compression level for all GRGs and node type `mem2_ssd1_v2_x64`, meaning all jobs were limited to 256GB of RAM. 64 cores were used for all jobs, and for a few of the larger chromosomes we explicitly increased the number of regions used (`--parts 192`) to decrease peak RAM usage. Table S4 shows the times, costs, and file sizes.

##### 4.3 UK Biobank PCA and GWAS

We ran PCA on all 490,541 individuals in the UK Biobank phased WGS dataset (field 30108), containing 137,116,837 variants after filtering out ultra-rare (allele count $< 20$) variants. We did not restrict to SNPs, remove related individuals, or perform any QC aside from removing ultra-rare variants. The PCA results available from the UK Biobank (field 22009) used fewer variants (147,604, about 900 times fewer than GRG used), and removed related individuals [3]. In Figure S4, you can see that the shapes of the top 3 PCs are similar between the two subsets of data, despite the differences in pre-processing, and the huge difference in the number of variants used.

We then looked at the usage of PCA for controlling population stratification in GWAS studies. Here we used a dataset restricted to unrelated white British individuals and removed all ultra-rare variants (allele count $< 20$), resulting in 338,117 individuals with 89,988,512 total variants. We compared the use of the top 20 PCs from three different methods: GRG-based PCA on all 22 autosomes for this dataset, GRG-based PCA on all 21 autosomes excluding the target chromosome (LOCO), and the PCA results from UK Biobank field 22009 that were generated after LD pruning (147,604 total variants used). We tested GWAS on two quantitative phenotypes - body mass index (BMI, field 21001) (Figure S9-S10) and standing height (field 50) (Figure S11-S12). We chose to examine chromosomes 3, 5, 6, 7, 8, 9, 10, for a mix of chromosomes that showed LD effect on the top 20 PCs (chromosomes 5, 6, 7, 8, 10) and ones that did not (chromosomes 3, 9).

First, we looked at the amount of correlation with the genotypes for the three different methods. We computed the correlation by running *grapp*'s linear regression (GWAS) and using each PC in turn as the phenotype ($Y$): e.g., GWAS on chromosome 1 against $Y = PC1$ for the "ALL" method, repeated for all PCs and all methods. Then we plotted the coefficient of determination ($R^2$) for each variant. Interestingly, for the chromosomes we tested there is a stronger genotype correlation with the LD-pruned PCs than with the LOCO PCs (Figure S5). This could be due to a number of factors, such as the variant subset being more likely to tag a large number of other variants (e.g., due to array design or the LD pruning itself), or just the magnitude difference in the number of variants being summarized by PCA (LOCO was generated from 600 times more variants than the LD-pruned PCs). The LOCO and LD-pruned PCs have fairly similar cumulative $R^2$ curves for all tested chromosomes, and deviate substantially from the $R^2$ curves for the PCs generated from all autosomes on most chromosomes (Figure S5). Chromosomes 6, 7, and 10 show high genotype/PC correlation within the first 10 PCs, whereas chromosomes 5 and 8 do not show high correlation until PCs 11-20.

Next we looked at the effect of PCA method on the p-values produced by GWAS. Here we used the top 20 PCs plus the individual's sex as covariates for GWAS against the body-mass index (BMI) and standing height (height) phenotypes. LOCO and LD pruned PCs used in GWAS produce very similar p-values for both BMI (Figure S10) and height (Figure S12). As expected, using PCs generated from all unpruned autosomes affect the p-value distribution, but only for certain chromosomes. Strong deviation in p-value distribution is visible for chromosomes 6 and 8 for BMI, and chromosome 6 for height, with some minor deviations present on other chromosomes (e.g., chromosome 8 for height). Since the LD pruning and leave-one-chromosome-out (LOCO) methods are consistent, it seems that these p-value deviations are caused by LD within the target chromosome. The only large, noticeable impact of this LD effect on the Manhattan plot of p-values is that of chromosome 8 for BMI, which has a broad spike of p-values between 1MBP and 2MBP on the LOCO and LD pruned GWAS plots.

### 5 Tables

Table S1: Constructing GRGs on UK Biobank 500,000 individuals WGS data: RAM usage and construction options.

| Chromosome | Options | Max RAM (MB) |
|---|---|---|
| 22 | `-j 64` | 94755 |
| 21 | `-j 64` | 96087 |
| 20 | `-j 64` | 141077 |
| 19 | `-j 64` | 141845 |
| 18 | `-j 64` | 155979 |
| 17 | `-j 64` | 176533 |
| 16 | `-j 64` | 185846 |
| 15 | `-j 64` | 157629 |
| 14 | `-j 64` | 168903 |
| 13 | `-j 64` | 190172 |
| 12 | `-j 64` | 248649 |
| 11 | `-j 64` | 224304 |
| 10 | `-j 64 --level1 --reduce 5` | 133210 |
| 9 | `-j 64` | 237728 |
| 8 | `-j 64 --level1 --reduce 5 --parts 192` | 134135 |
| 7 | `-j 64 --level1 --reduce 5 --parts 192` | 171847 |
| 6 | `-j 64 --level1 --reduce 5 --parts 192` | 195282 |
| 5 | `-j 64 --level1 --reduce 5 --parts 192` | 180092 |
| 4 | `-j 64 --level1 --reduce 5 --parts 192` | 190398 |
| 3 | `-j 64 --level1 --reduce 5 --parts 192` | 195755 |
| 2 | `-j 64 --level1 --reduce 5 --parts 192` | 247619 |
| 1 | `-j 64 --level1 --reduce 5 --parts 192` | 218127 |

The data is from the UK Biobank logs, which reports RAM usage periodically. Increasing the `-p` option (default: 100) reduces RAM usage by splitting the chromosome into more parts. The `--level1` option builds smaller trees, which also reduces RAM (but `--reduce 5` makes sure we still optimize the graph size afterwards).

Table S2: Comparison of PGEN and GRG construction on UK Biobank chromosome 22, phased WGS data.

| Chrom. | Format | Resulting file Size (GB) | Threads | From | Minutes Elapsed | Minutes (CPU) |
|---|---|---|---|---|---|---|
| 22 | PGEN | 17.40 | 2 | .vcf.gz | 1247.62 | 2495.24 |
| 22 | PGEN | 17.40 | 70 | .vcf.gz | 335.40 | 23478.00 |
| 22 | GRG | 1.98 | 70 | .vcf.gz | 100.72 | 7050.40 |
| 22 | GRG | 2.03 | 70 | IGD | 36.60 | 2562.00 |

Table S3: Comparison of PGEN and GRG construction on simulated data.

| Dataset | Diploids | Format | Resulting file size (GB) | Threads | From | Minutes Elapsed |
|---|---|---|---|---|---|---|
| Simulated | 100000 | PGEN | 13.00 | 25 | .vcf.gz | 121.00 |
| Simulated | 100000 | GRG | 1.10 | 25 | .vcf.gz | 44.37 |
| Simulated | 100000 | GRG (fast) | 1.40 | 25 | .vcf.gz | 34.00 |
| Simulated | 100000 | GRG | 1.10 | 25 | IGD | 15.19 |
| Simulated | 100000 | GRG (fast) | 1.40 | 25 | IGD | 6.55 |
| Simulated | 200000 | PGEN | 25.00 | 25 | .vcf.gz | 265.62 |
| Simulated | 200000 | GRG | 1.50 | 25 | .vcf.gz | 91.60 |
| Simulated | 200000 | GRG (fast) | 2.00 | 25 | .vcf.gz | 75.33 |
| Simulated | 200000 | GRG | 1.50 | 25 | IGD | 24.93 |
| Simulated | 200000 | GRG (fast) | 2.00 | 25 | IGD | 10.73 |
| Simulated | 500000 | PGEN | 64.00 | 25 | .vcf.gz | 790.88 |
| Simulated | 500000 | GRG | 2.00 | 25 | .vcf.gz | 251.38 |
| Simulated | 500000 | GRG (fast) | 3.90 | 25 | .vcf.gz | 218.48 |
| Simulated | 500000 | GRG | 2.00 | 25 | IGD | 51.11 |
| Simulated | 500000 | GRG (fast) | 3.90 | 25 | IGD | 22.62 |
| Simulated | 1000000 | PGEN | 127.00 | 25 | .vcf.gz | 1810.03 |
| Simulated | 1000000 | GRG | 2.70 | 25 | .vcf.gz | 564.40 |
| Simulated | 1000000 | GRG (fast) | 7.00 | 25 | .vcf.gz | 502.85 |
| Simulated | 1000000 | GRG | 2.70 | 25 | IGD | 91.78 |
| Simulated | 1000000 | GRG (fast) | 7.00 | 25 | IGD | 42.19 |

Table S4: Constructing GRGs on UK Biobank 500,000 individuals WGS data.

| Chromosome | VCF Size | GRG Size[1] | GRG Cost[2] | GRG Hours | Nodes | Edges |
|---|---|---|---|---|---|---|
| 22 | 43.29 | 2.03 | 0.35 | 0.61 | 44890619 | 597707867 |
| 21 | 41.45 | 1.86 | 1.61 | 0.68 | 40826747 | 554222453 |
| 20 | 69.90 | 3.00 | 2.66 | 1.01 | 67845552 | 843463506 |
| 19 | 70.66 | 3.27 | 0.55 | 1.10 | 74047918 | 941920073 |
| 18 | 85.84 | 3.46 | 2.47 | 1.15 | 79106026 | 951091613 |
| 17 | 87.46 | 3.93 | 3.39 | 1.60 | 89886895 | 1106083118 |
| 16 | 97.92 | 4.20 | 3.04 | 1.40 | 94270109 | 1163413068 |
| 15 | 89.27 | 3.74 | 3.26 | 1.29 | 85292948 | 1045908195 |
| 14 | 99.30 | 3.86 | 3.18 | 1.55 | 86948014 | 1057329816 |
| 13 | 110.10 | 4.19 | 3.79 | 1.56 | 95167526 | 1141052543 |
| 12 | 146.99 | 5.68 | 4.92 | 2.15 | 128439887 | 1542757615 |
| 11 | 151.24 | 5.60 | 3.85 | 1.86 | 125700064 | 1504178301 |
| 10 | 152.39 | 6.18 | 4.70 | 2.25 | 110307100 | 2074318574 |
| 9 | 135.16 | 5.41 | 3.98 | 1.89 | 120632165 | 1501775324 |
| 8 | 168.67 | 6.53 | 4.21 | 2.03 | 114014670 | 2165223710 |
| 7 | 180.09 | 7.24 | 4.68 | 2.26 | 127297908 | 2429052011 |
| 6 | 195.08 | 7.24 | 4.96 | 2.38 | 127041754 | 2406642972 |
| 5 | 198.89 | 7.57 | 5.36 | 2.57 | 133278665 | 2495969487 |
| 4 | 217.73 | 8.16 | 6.22 | 2.99 | 143837344 | 2687908635 |
| 3 | 219.98 | 8.43 | 6.10 | 2.94 | 149509140 | 2765478526 |
| 2 | 263.30 | 10.40 | 7.38 | 3.57 | 184784321 | 3285887694 |
| 1 | 243.72 | 9.95 | 7.05 | 3.39 | 178205219 | 3285887694 |

[1]Sizes are in Gigabytes, cost is in GBP. 64 threads were used for each chromosome.

[2]Costs are for construction from IGD on a mixture of spot and on-demand instances; larger chromosomes used on-demand jobs.

Table S5: Comparing PCA methods on unfiltered simulated data.

| Method | Threads[1] | Minutes | Individuals | RAM (GB)[2] |
|---|---|---|---|---|
| GRG | 1 | 6.57 | 100000 | 1.97 |
| GRG | 1 | 14.25 | 500000 | 3.28 |
| GRG | 1 | 15.50 | 1000000 | 4.21 |
| GRG ProPCA | 1 | 6.22 | 100000 | 13.88 |
| GRG ProPCA | 1 | 12.64 | 500000 | 21.87 |
| GRG ProPCA | 1 | 17.68 | 1000000 | 26.14 |
| FlashPCA2 | 1 | 1359.42 | 100000 | 99.52 |
| FlashPCA2 | 1 | 12826.00 | 500000 | 103.09 |
| FlashPCA2 | 1 | Did not finish | 1000000 | Did not finish |
| plink PGEN | 25 | 340.03 | 100000 | 66.32 |
| plink PGEN | 25 | 2348.50 | 500000 | 116.99 |
| plink PGEN | 25 | 7630.58 | 1000000 | 116.51 |

[1]Threads is the number of threads *utilized.*

[2]Jobs were given a maximum of 128GB RAM and 240 hours.

Table S6: Comparing PCA methods on simulated data filtered so that allele frequency $\geq 0.05$.

| Method | Threads[1] | Minutes | Individuals | RAM (Gigabytes)[2] |
|---|---|---|---|---|
| GRG | 1 | 6.57 | 100000 | 0.91 |
| GRG | 1 | 15.93 | 500000 | 1.74 |
| GRG | 1 | 16.60 | 1000000 | 2.41 |
| GRG ProPCA | 1 | 5.00 | 100000 | 3.81 |
| GRG ProPCA | 1 | 9.49 | 500000 | 7.06 |
| GRG ProPCA | 1 | 14.59 | 1000000 | 9.21 |
| FlashPCA2 | 1 | 45.17 | 100000 | 100.02 |
| FlashPCA2 | 1 | 228.60 | 500000 | 100.13 |
| FlashPCA2 | 1 | 343.13 | 1000000 | 100.24 |
| plink PGEN | 25 | 16.65 | 100000 | 18.39 |
| plink PGEN | 25 | 83.27 | 500000 | 85.73 |
| plink PGEN | 25 | 174.67 | 1000000 | 114.69 |

[1]Threads is the number of threads *utilized.*

[2]Jobs were given a maximum of 128GB RAM and 240 hours.

Table S7: Comparison of XSI, Savvy, and GRG on UK Biobank 200,000 individual WGS data.

| Chromosome | Format[1] | Size (GB) | Hours | Threads | CPU Hours |
|---|---|---|---|---|---|
| 13 | GRG v1 | 5.91 | 5.00 | 70 | 350.08 |
| 13 | GRG v2 | 2.03 | 0.45 | 70 | 31.50 |
| 13 | Savvy | 3.34 | 13.80 | 1 | 13.80 |
| 13 | XSI | 2.76 | 21.96 | 1 | 21.96 |
| 22 | GRG v1 | 2.40 | 4.57 | 70 | 320.07 |
| 22 | GRG v2 | 0.90 | 0.16 | 70 | 11.20 |
| 22 | Savvy | 1.35 | 5.07 | 1 | 5.07 |
| 22 | XSI | 1.12 | 8.03 | 1 | 8.03 |

[1]GRG was constructed from IGD, XSI and Savvy were constructed from BCF. We used the default block sizes for both XSI and Savvy in this experiment. See Figure S13 for a more detailed format comparison.

Table S8: GRG load time and matvec time on GRGs constructed from simulated data. Average of 10 replicates.

| Individuals | Noise[1] | File Size (GB) | Load Time (Sec) | Matvec Time (Sec) |
|---|---|---|---|---|
| 100,000 | No | 1.2 | 2.31 | 3.32 |
| 100,000 | Yes | 3.9 | 3.57 | 14.56 |
| 200,000 | No | 1.5 | 2.31 | 4.45 |
| 200,000 | Yes | 6.1 | 3.94 | 23.16 |

[1]Noise was added to the simulated data in the same way as described in Section 3.

### 6 Figures

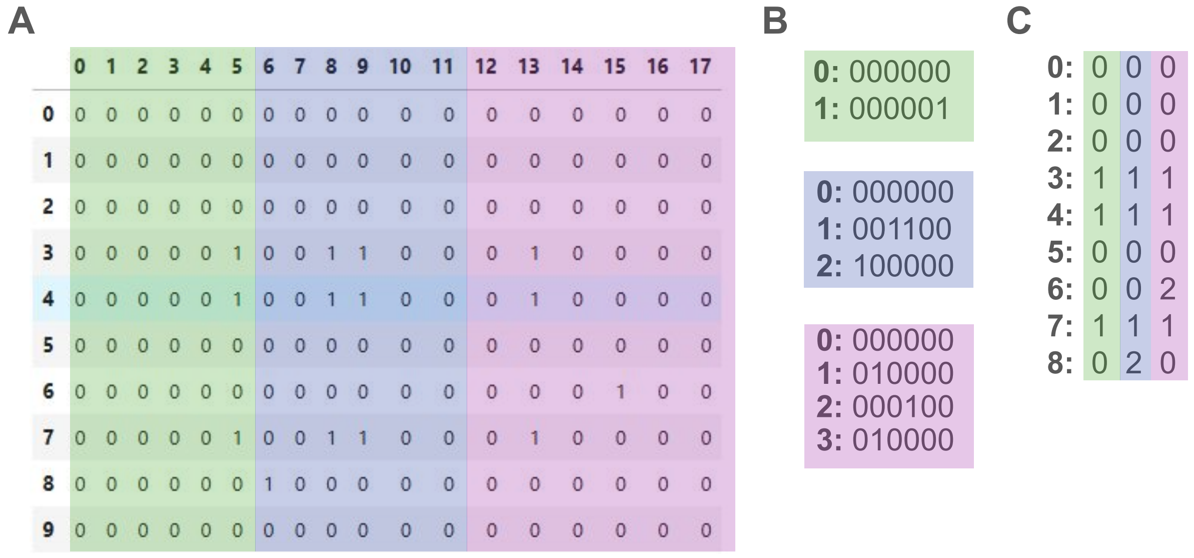

Figure S1: **Diagram of haplotype segment representation. A.** A genotype matrix with 10 haplotypes (y-axis) and 18 variants (x-axis). Three regions of 6 variants each are highlighted in green, blue, and purple. **B.** The unique haplotype segments for each of the 6 regions, numbered (on the left) by the haplotype segment ID. **C.** The original 10 haplotypes, represented by a vector of their haplotype segment IDs instead of directly representing their haplotypes.

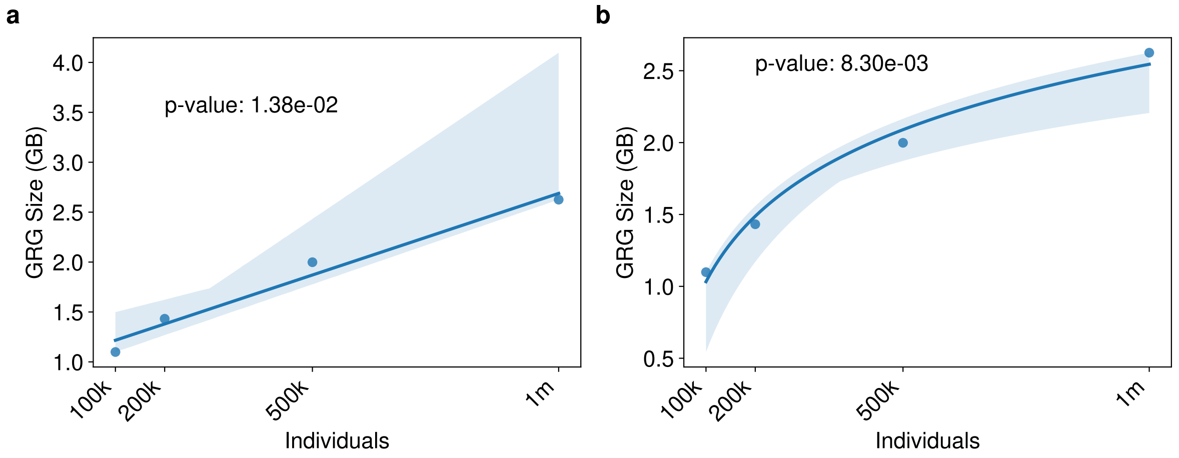

Figure S2: **GRG constructed file sizes for simulated data. a.** Linear regression of sample size (x-axis) against GRG file size (y-axis), with p-value. **b.** Same linear regression, except against $log(Individuals)$.

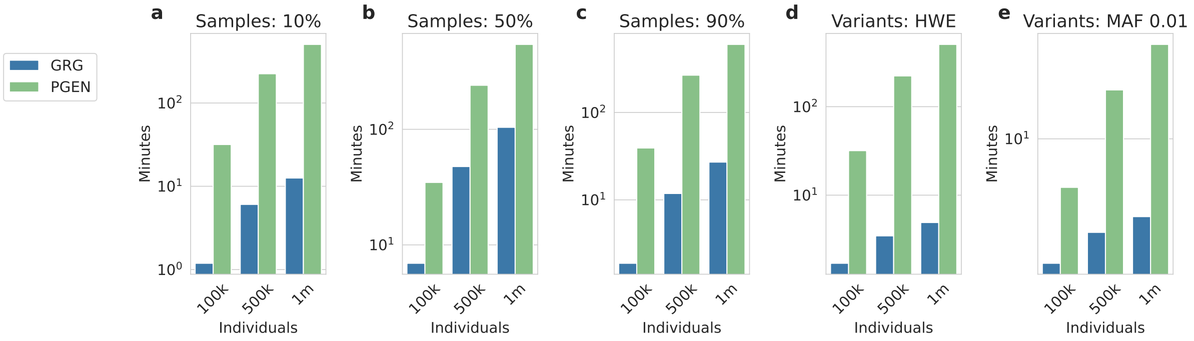

Figure S3: **Comparison of dataset filtering performance between *PLINK2* and *grapp*. a.** Keep the first 10% of samples. **b.** Keep the first 50% of samples. **c.** Keep the first 90% of samples. **d.** Keep only variants with Hardy-Weinberg p-value greater than $1 \times 10^{-6}$. **e.** Keep only variants with minor allele frequency $\geq 0.01$. All tests were run on simulated data, with number of individuals on the y-axis. Time (x-axis) is log scale.

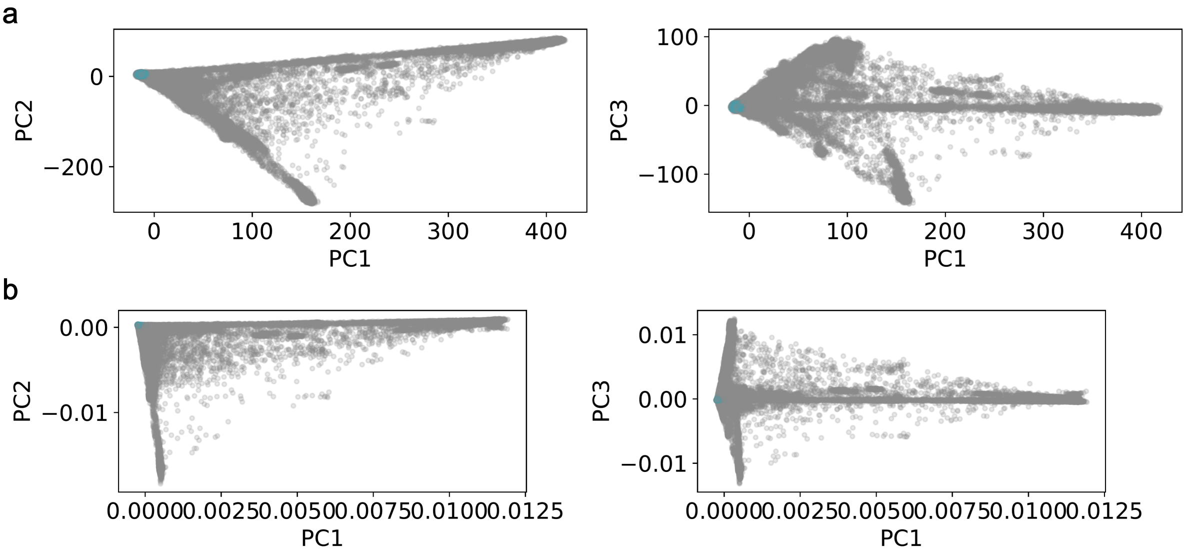

Figure S4: **UKB PCs for pruned, sparse data vs WGS. a.** PC1,2 and PC1,3 plots of PCA results provided by UK Biobank (field 22009), which were generated from 147,604 variants of 407,219 unrelated samples [3]. **b.** PC1,2 and PC1,3 plots of GRG PCA results, which were generated from 137,116,837 variants of 490,541 individuals.

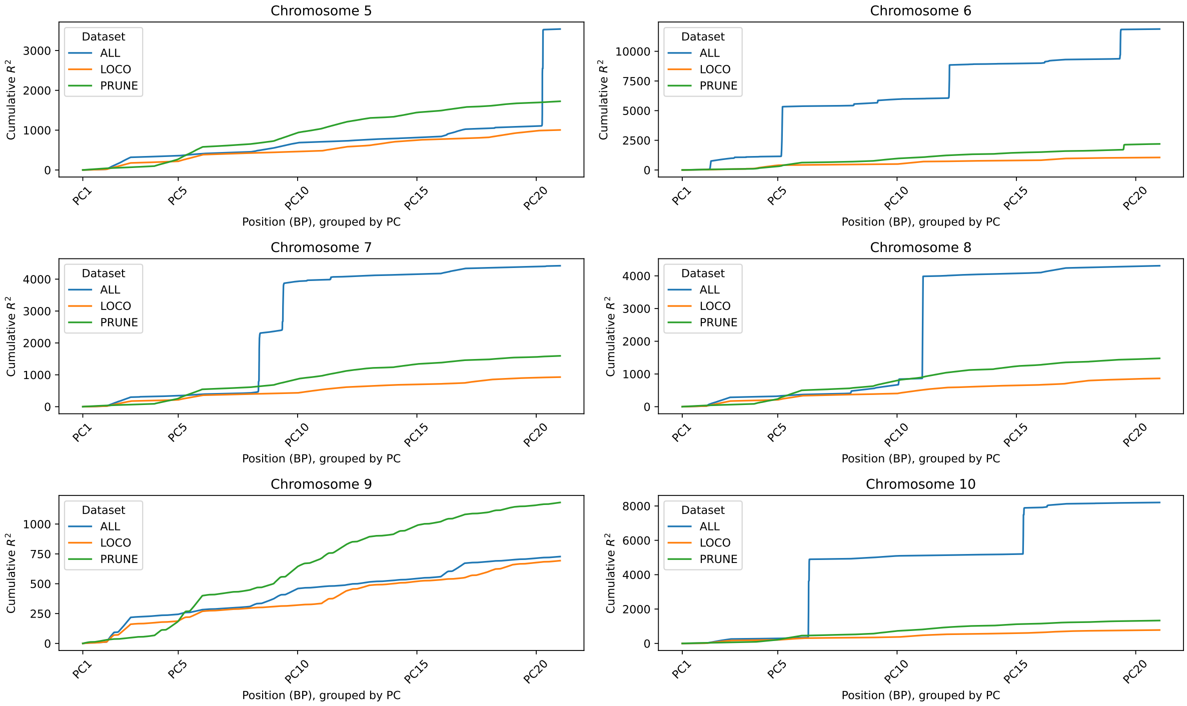

Figure S5: **PC-genotype correlation for different PCA methods.** The y-axis is the cumulative $R^2$ from comparing the PC (x-axis) with the genotype of the chromosome. The x-axis shows the cumulative correlation by position within each PC, so 0 on the x-axis is base-pair position 0 compared with PC1, and the last position is the last base-pair position compared with PC20. The cumulative $R^2$ is computed from every variant in the genotype (UKB white British, variants with allele count $\geq 20$), but only every $500^{th}$ point is plotted.

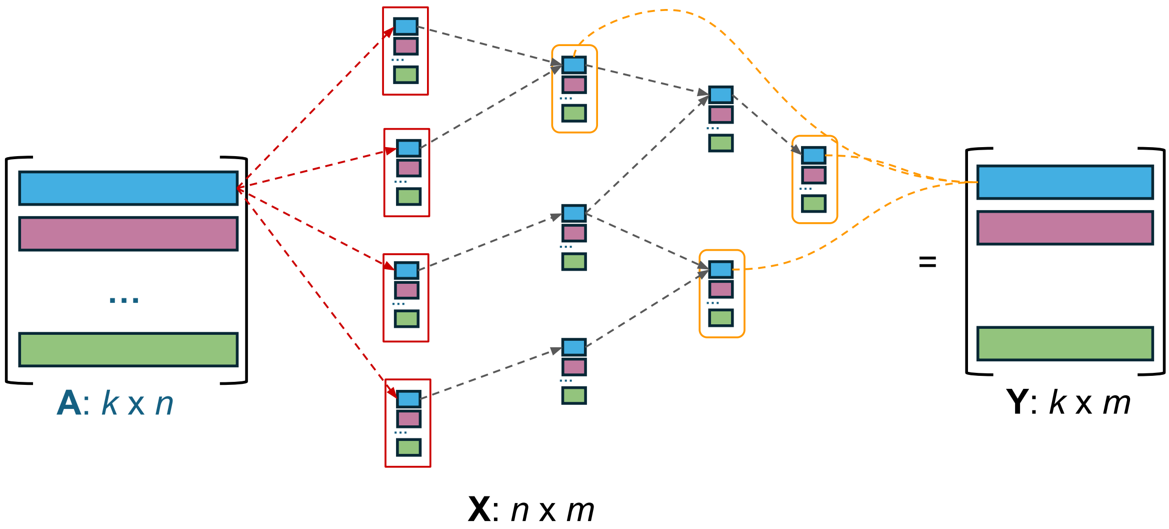

Figure S6: **Diagram of matrix multiplication with GRG.** Each colored row in the input (output) matrix **A** (**B**) propagates to/from the identically colored value holders (small rectangles) for each node. The GRG nodes and edges are not illustrated directly, but instead the dashed edges show how values are propagated between the values associated with each node. Only the first (blue) row is shown; the other colors will similarly propagate to the value holders of the same color. The red edges/boxes correspond to *sample nodes*. The orange edges/boxes correspond to *mutation nodes*.

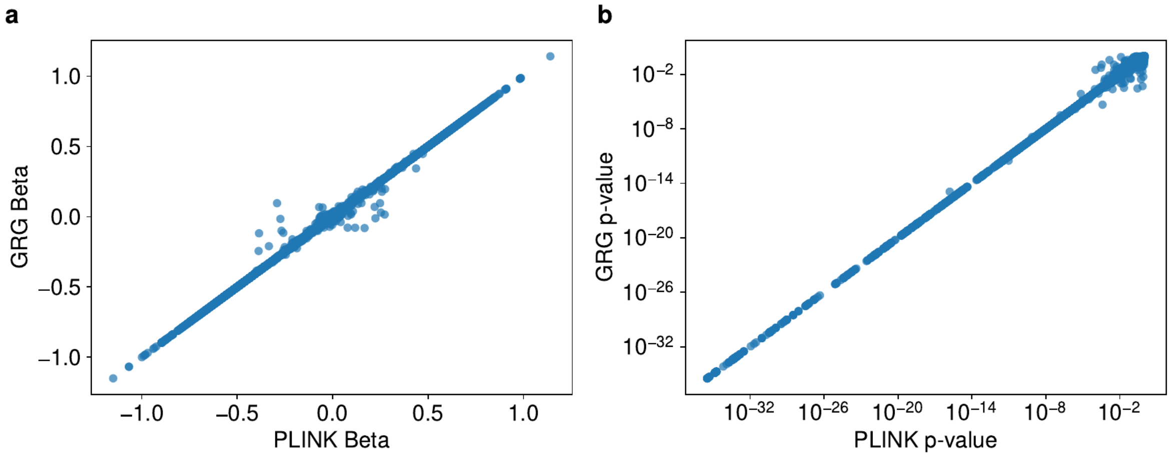

Figure S7: **Comparison of $\beta$ and p-values between *PLINK2* and *grapp*.** The p-values are plotted on a log-log scale so it is easier to see the outlier (more likely to be significant) values. Test performed on simulated data with 25,000 diploid individuals and a simulated phenotype, with the top 10 PCs used as covariates.

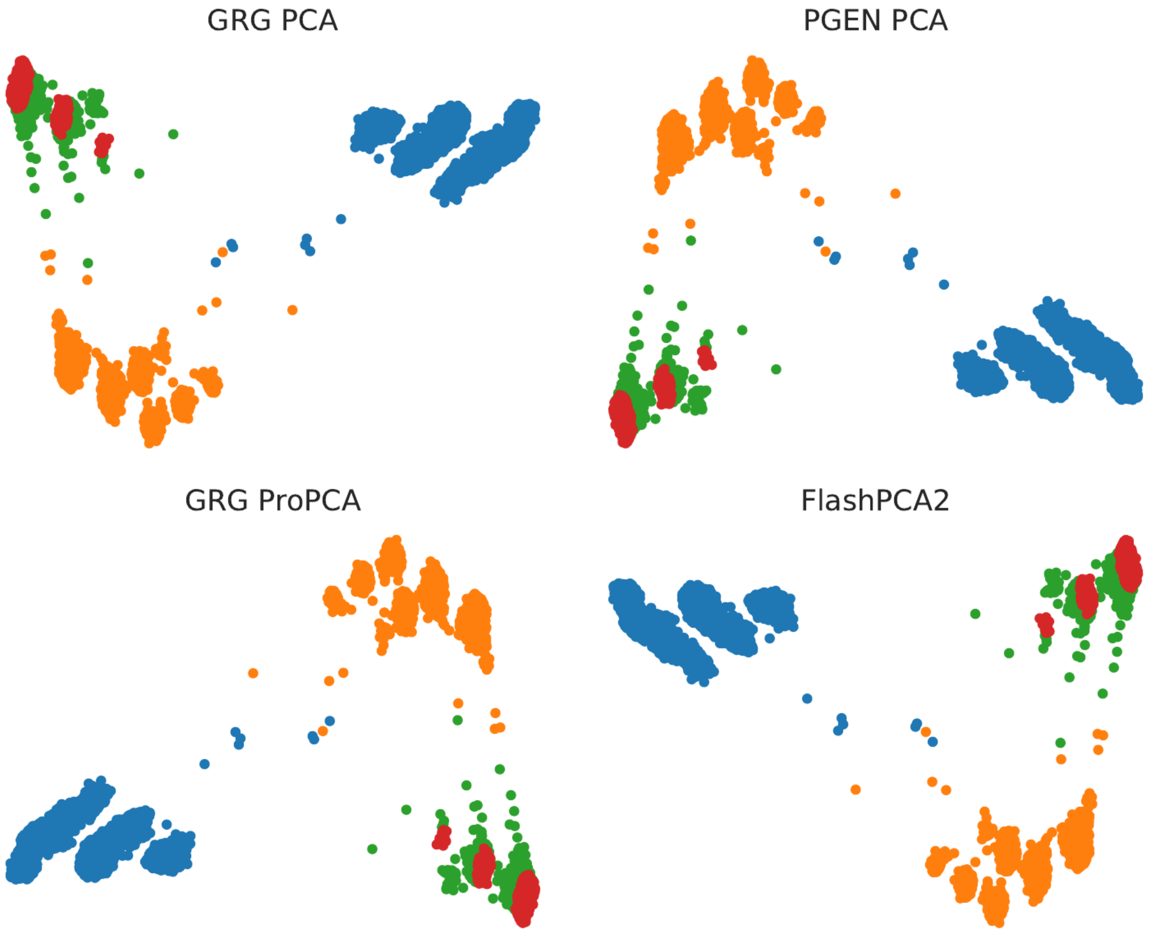

Figure S8: **PC1 vs. PC2 on simulated OutOfAfrica_4J17 dataset.** Comparison between GRG (eigen decomposition method), GRG ProPCA implementation, plink2's PGEN-based PCA, and Flash-PCA2. Rotations about the $x$ or $y$ axes can result from slight numerical differences during PCA solve.

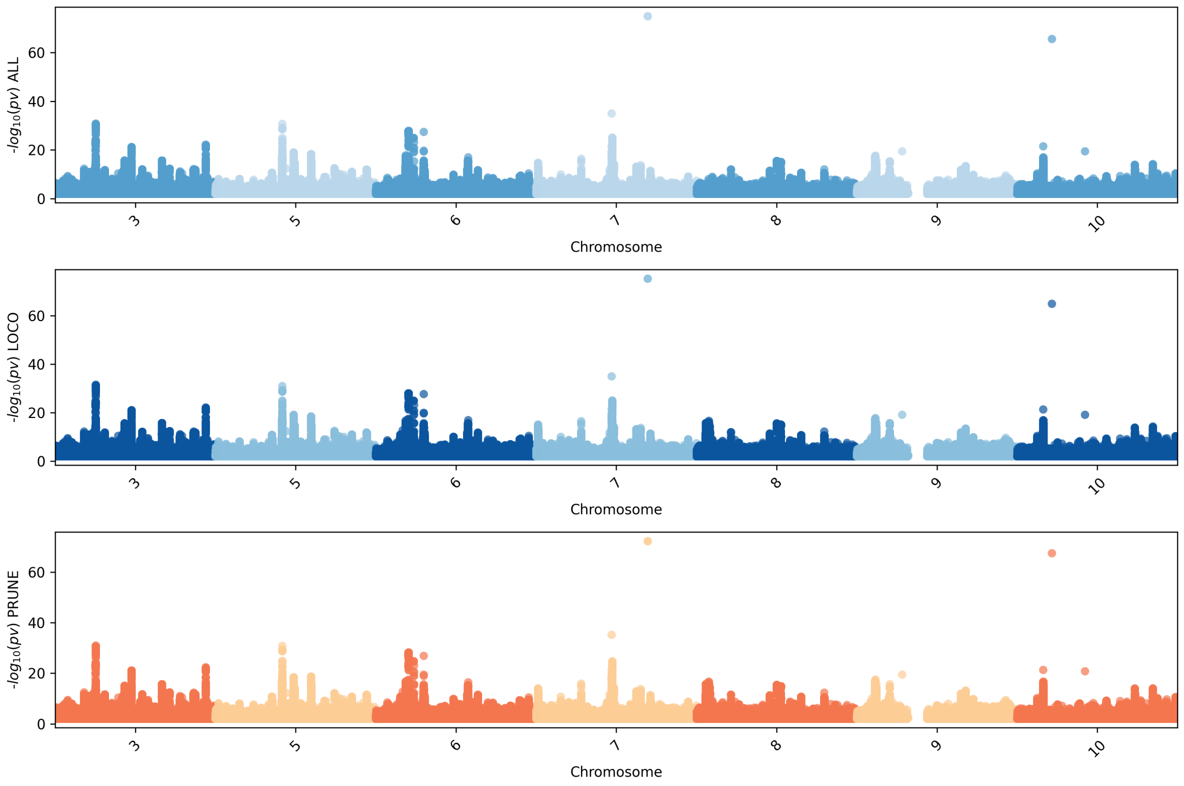

Figure S9: **Manhattan plot for BMI GWAS with different PCA methods.** GWAS p-values for chromosomes 3, 5-10, with position on x-axis and $-log_{10}(p\text{-}value)$ on y-axis. The three rows use different PCA methods to generate PCs for covariates: PCA on all unpruned autosomes (ALL, top), PCA with leave-one-chromosome-out (LOCO, middle), and LD pruned dataset for PCA (PRUNE, bottom).

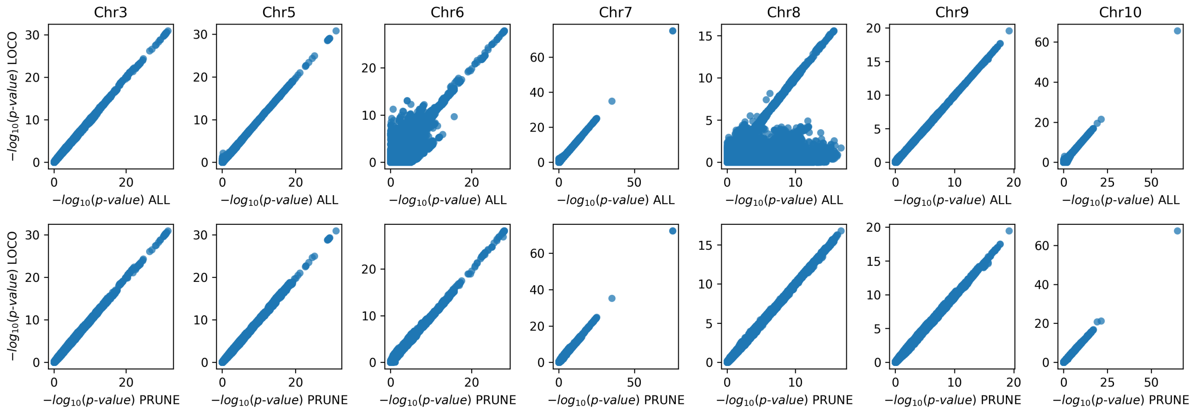

Figure S10: **Comparing p-values for BMI GWAS runs with different PCA methods.** The upper row plots $-log_{10}(p\text{-}value)$ for LOCO PCs (y-axis) vs. ALL PCs (x-axis). The bottom row compares LOCO PCs (y-axis) vs. LD-pruned PCs (UK Biobank field 22009).

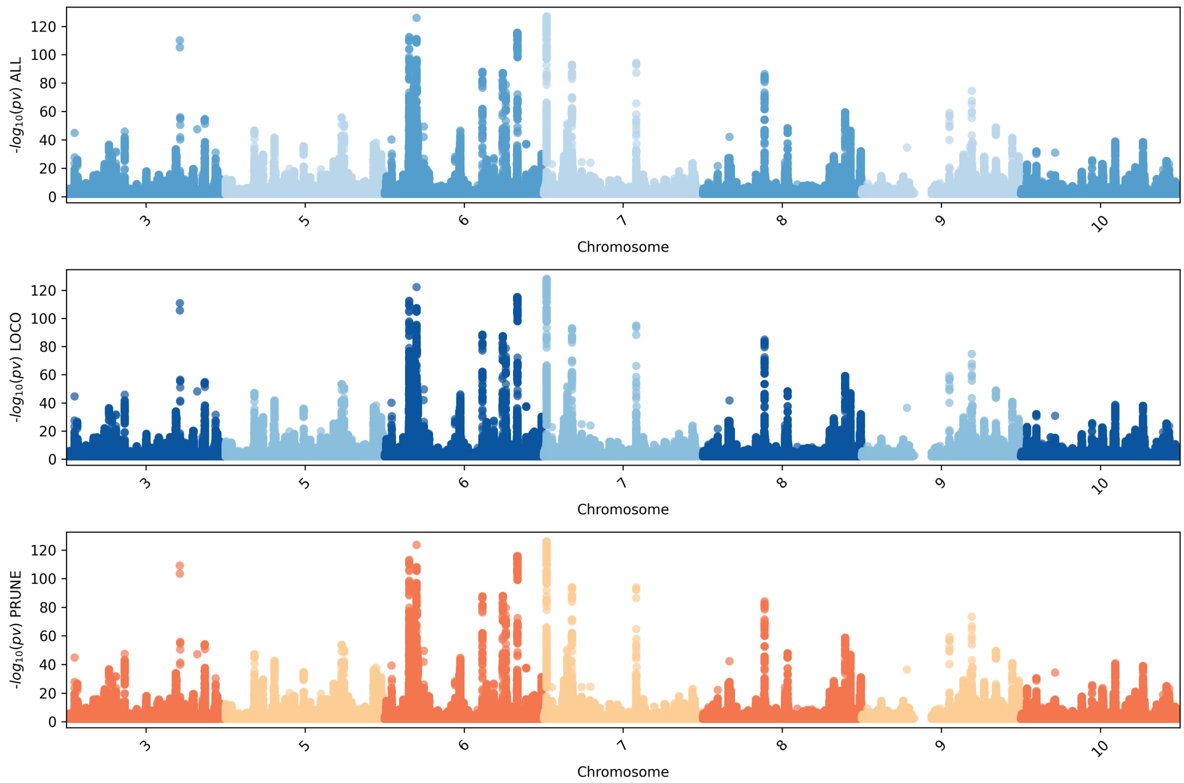

Figure S11: **Manhattan plot for "standing height" GWAS with different PCA methods.** GWAS p-values for chromosomes 3, 5-10, with position on x-axis and $-log_{10}(p\text{-}value)$ on y-axis. The three rows use different PCA methods to generate PCs for covariates: PCA on all unpruned autosomes (ALL, top), PCA with leave-one-chromosome-out (LOCO, middle), and LD pruned dataset for PCA (PRUNE, bottom).

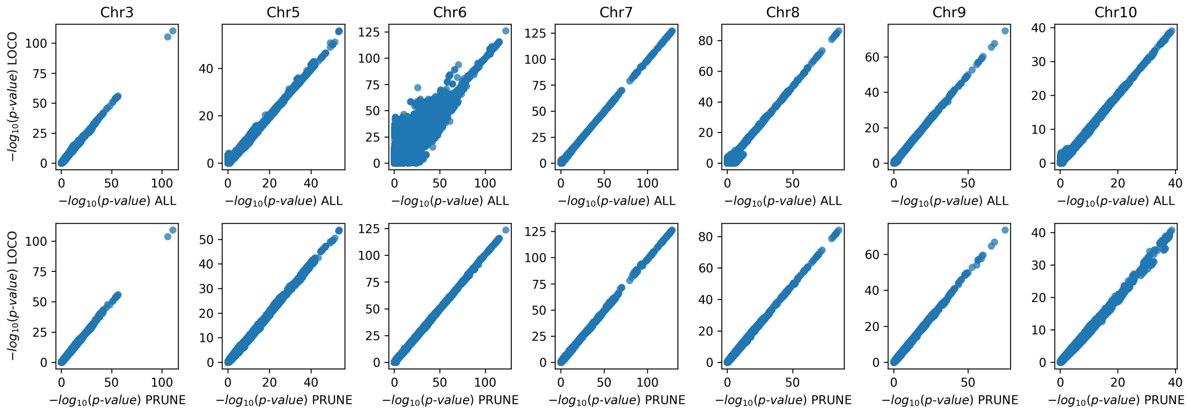

Figure S12: **Comparing p-values for "standing height" GWAS runs with different PCA methods.** The upper row plots $-log_{10}(p\text{-}value)$ for LOCO PCs (y-axis) vs. ALL PCs (x-axis). The bottom row compares LOCO PCs (y-axis) vs. LD-pruned PCs (UK Biobank field 22009).

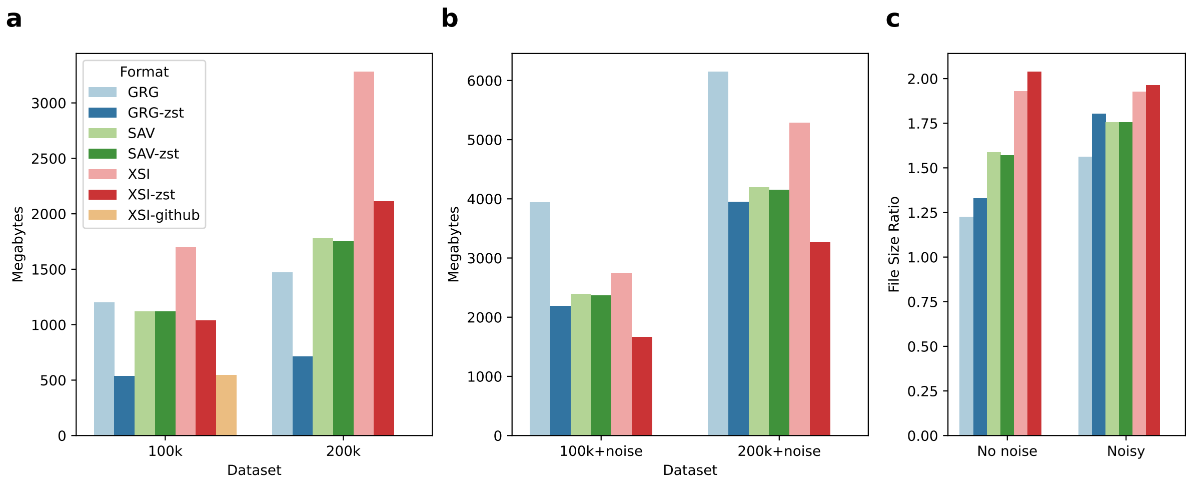

Figure S13: **Compression by file format.** Data is simulated 100,000 or 200,000 diploid individuals, with or without noise. **a.** File size on data without noise. **b.** File size on data with noise. **c.** Ratio of the file size for 200,000 individuals to the file size for 100,000 individuals. This represents a scaling factor for the file format as the sample size increases. The "-zst" suffix indicates the file was compressed using the zstandard compression tool, after creating it with the corresponding tool (GRG, Savvy, XSI). Results use Savvy v1.5.5 and XSI v4.0.0. "XSI-github" is the untagged latest GitHub version, which supports variable-sized blocks, and was run with a large block size and built-in zstd compression. See Section 3.1 for full details.

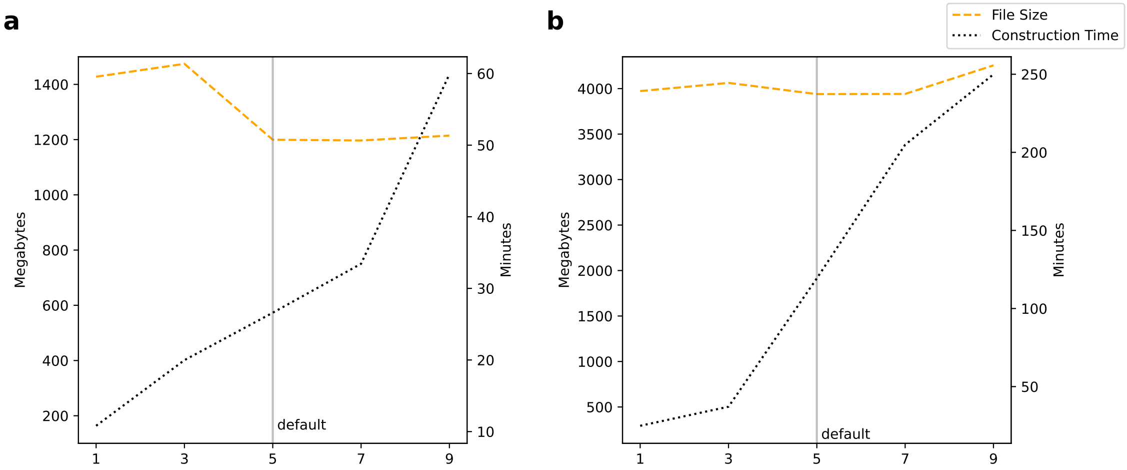

Figure S14: **GRG compression level.** Data is simulated 100,000 or 200,000 diploid individuals, with or without noise. Compression levels 1, 3, 5, 7, and 9 were tested. **a.** Data without noise. Left y-axis (orange line) is file size, right y-axis (black line) is construct time. **b.** Same as panel **a** except for noisy data.

[4] Yann Collet. Rfc 8878: Zstandard compression and the'application/zstd'media type, 2021.

[5] Drew DeHaas and Xinzhu Wei. Igd: a simple, efficient genotype data format. *Bioinformatics Advances*, 5(1):vbaf205, 08 2025. ISSN 2635-0041. `doi:10.1093/bioadv/vbaf205`. `https://academic.oup.com/bioinformaticsadvances/article-pdf/5/1/vbaf205/64140915/vbaf205.pdf`. URL `https://doi.org/10.1093/bioadv/vbaf205`.

[6] Drew DeHaas, Ziqing Pan, and Xinzhu Wei. Enabling efficient analysis of biobank-scale data with genotype representation graphs. *Nature computational science*, 5(2):112–124, 2025.

[7] Jonathon LeFaive, Albert V Smith, Hyun Min Kang, and Goncalo Abecasis. Sparse allele vectors and the savvy software suite. *Bioinformatics*, 37(22):4248–4250, 2021.

[8] Richard B Lehoucq, Danny C Sorensen, and Chao Yang. *ARPACK users' guide: solution of large-scale eigenvalue problems with implicitly restarted Arnoldi methods*. SIAM, 1998.

[9] John; Lemire Daniel Rupp, Christoph; Lees-Miller. libvbyte - fast c library for 32bit and 64bit integer compression. `https://github.com/cruppstahl/libvbyte`, 2025.

[10] Rick Wertenbroek, Simone Rubinacci, Ioannis Xenarios, Yann Thoma, and Olivier Delaneau. Xsi—a genotype compression tool for compressive genomics in large biobanks. *Bioinformatics*, 38(15):3778–3784, 2022.
